# Membrane Curvature Activates Src kinase and Promotes Metastatic Cancer Cell Survival

**DOI:** 10.64898/2026.03.03.709279

**Authors:** Wei Zhang, He You, Xinzhi Zou, Chih-Hao Lu, Xingyuan Zhang, Anna Amine, Zeinab Jahed, Michael Z. Lin, Bianxiao Cui

## Abstract

Src family kinases (SFKs) play key roles in cancer metastasis. While SFKs are classically regulated by cell adhesions and transmembrane receptors, how they become activated following tumor cell detachment remains unclear. Here, we report curvature-induced kinase activation (CIKA), a distinct mechanism through which plasma membrane curvature directly promotes Src activation. Mechanistically, membrane curvature promotes the oligomerization of TOCA-family curvature-sensing proteins, inducing local biomolecular condensation. These condensates recruit Src, stabilize its open conformation, and exclude the negative regulator Csk, converting curved membrane domains into discrete kinase activation hubs. Disruption of CIKA using TOCA mutants inhibits curvature-induced Src activation and selectively impairs the viability of detached but not adherent cells. Functionally, curvature-induced Src activation promotes anchorage-independent survival, and its disruption suppresses metastatic colonization in xenograft mouse models. These findings reveal membrane curvature as a biophysical activator of Src and suggest CIKA inhibition as a potential therapeutic strategy to target metastatic cancer cells.

## Main

Src family kinases (SFKs) are key regulators of cellular signaling. Src was the first oncogene and the first tyrosine kinase identified, establishing the paradigm of non-receptor tyrosine kinases^1–4^. Aberrant Src activation is frequent in solid tumors and drives malignant progression^5–8^. In adherent cells, cell–matrix and cell–cell adhesions play central roles in Src activation, where adhesion receptors organize signaling complexes that locally recruit and activate Src^9–11^. During metastasis, however, tumor cells detach from surrounding tissues. In order to survive, they must evade anoikis, a form of cell death triggered by loss of adhesion^12,13^. In many tumor types, Src activity persists and often increases in detached cells, which is strongly associated with anoikis resistance^14–18^. Yet how Src becomes activated in the absence of adhesion remains poorly understood.

Metastatic cells are often softer than their non-metastatic counterparts and exhibit reduced plasma-membrane tension that favors membrane deformation^19–23^. Upon detachment, relaxation of cytoskeletal stress further reduces membrane tension^24,25^. Consistent with this, detached metastatic cells display highly ruffled surfaces that contain abundant curved membrane structures such as blebs, protrusions, and invaginations, which contribute to malignancy^21,26–30^. Beyond a mere geometric feature, membrane curvature has emerged as a biophysical signal^31^, regulating diverse cellular processes, including membrane trafficking^32,33^, adhesion^34,35^, GTPase signaling^36–39^, and transmembrane signaling^40–43^. SFKs are anchored to the inner leaflet of the plasma membrane, making them susceptible to regulation by membrane geometry. We therefore hypothesized that membrane curvature could influence Src activation, particularly during metastatic dissemination.

Src activity is canonically regulated by conformational changes^44^. In the inactive state, the SH2 and SH3 domains engage intramolecular ligands, locking Src in a closed, autoinhibited conformation^45,46^. Binding of external partners releases this autoinhibition, allowing autophosphorylation at Y419 within the catalytic domain to achieve full kinase activity^44,47,48^. Recent studies have shown that phase-separated condensates can locally concentrate signaling molecules to facilitate robust and selective kinase activation^49,50^. However, condensation-based regulation of SFKs has not yet been observed in cells. Membrane curvature can potentially provide a platform for condensate formation, as many curvature-sensing proteins cluster at curved membranes and contain SH3 domains, a class of interaction modules that can mediate phase condensation^31,51–53^.

In this study, we used nanofabricated substrates to impose defined nanoscale curvature on the plasma membrane^54^. We identified curvature-induced kinase activation (CIKA) as a mechanism by which membrane curvature promotes Src activation and downstream phosphoinositide-3-kinase/protein kinase B (PI3K/Akt) signaling. Positively curved regions of the plasma membrane trigger liquid-like phase condensation by driving oligomerization of TOCA-family curvature-sensing proteins. Through multivalent interactions between SH3 domains and proline-rich motifs (PRMs), these proteins assemble adaptor proteins into condensates that concentrate Src. Within these condensates, Src adopts an open conformation and undergoes enhanced autophosphorylation at Y419, while the negative regulator C-terminal Src kinase (Csk) is excluded, establishing spatially restricted kinase-activation hubs. Curvature-induced Src activation promotes resistance to anoikis, a critical barrier to metastasis. Disrupting it using a dominant-negative FBP17(ΔSH3) construct selectively impairs anchorage-independent cell survival, reduces tumor-colony formation, and suppresses metastatic colonization in xenograft models. Together, these findings reveal curvature-induced Src activation as the mechanism linking membrane shape to Src activation and oncogenic signaling.

## Results

### Plasma membrane curvature enriches active Src family kinases

To investigate how membrane curvature influences Src activation, we used a nanofabricated platform that precisely controls local membrane geometry^54^. The platform consists of vertical silicon-dioxide nanopillars (200-500 nm diameter, 1.5-μm height) or nanobars (200-nm width and 1.2-μm height) engineered by electron-beam lithography (**Fig. 1a**). When cultured on these substrates, cells wrap their plasma membranes around nanostructures. Nanopillars generate cylindrical curvature and nanobar ends induce half-cylindrical curvature. Nanobars contain both curved (ends) and flat (sides) regions for direct comparison. Each substrate also includes flat regions serving as internal controls.

**Fig. 1:**
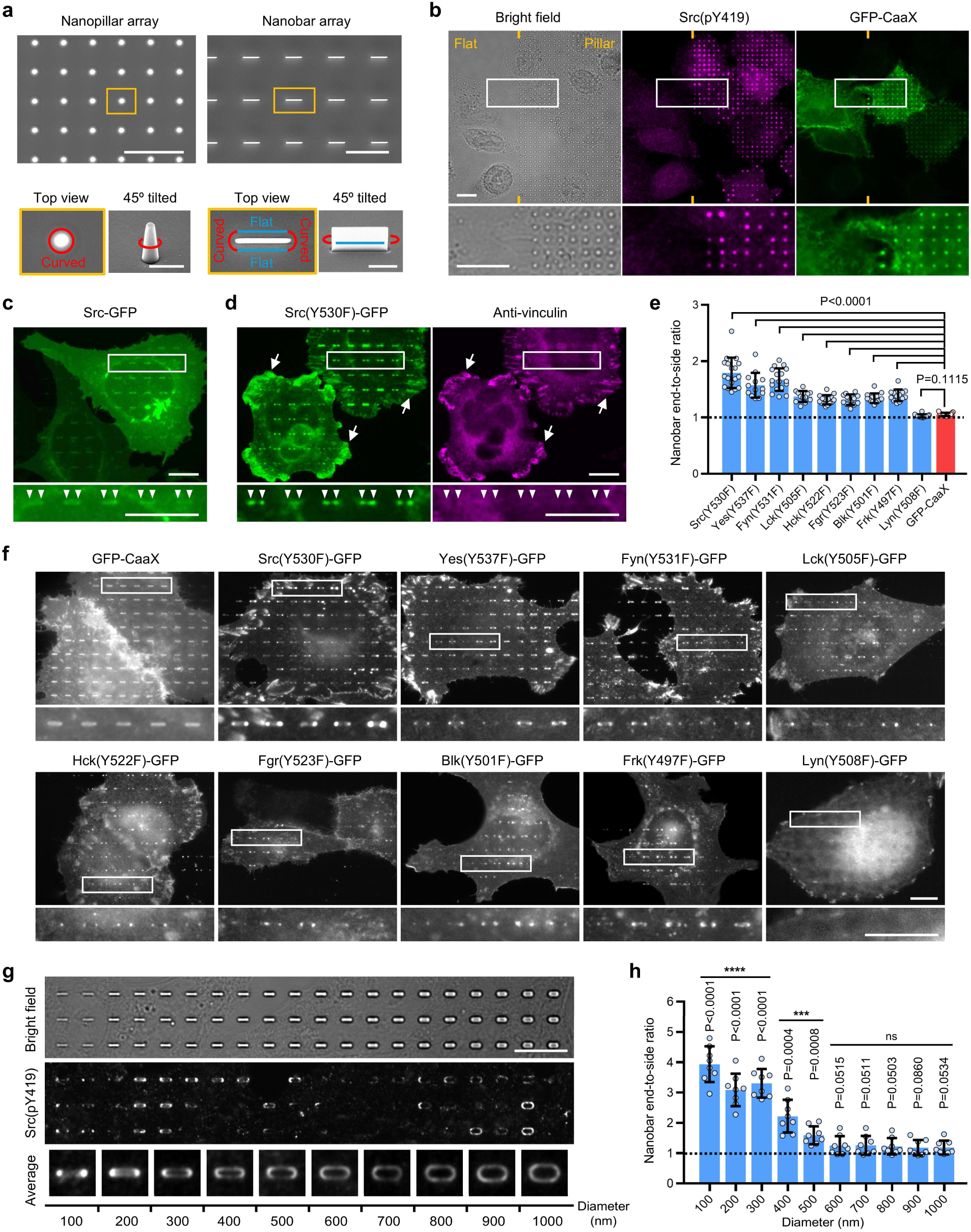
Plasma membrane curvature enriches active Src family kinases. **a**, Scanning electron microscopy (SEM) images of nanopillar and nanobar arrays. Schematics illustrate the local membrane curvature induced by these nanostructures. Scale bars: 5 µm (top), 1 µm (bottom). **b**, Immunostaining of active Src (Src(pY419)) shows selective enrichment at a subset of nanopillars, despite uniform membrane wrapping marked by GFP–CaaX. Scale bars, 10 µm. **c**, Wild-type Src–GFP shows a uniform membrane distribution without detectable enrichment at nanobar ends (arrows). Scale bars, 10 µm. **d**, Open-conformation Src mutant Src(Y530F)–GFP is strongly enriched at nanobar ends (arrowheads) and focal adhesions (arrows). Co-staining with vinculin confirms that the curvature enrichment is independent of focal adhesions. Scale bars, 10 µm. **e,** Quantification of nanobar end-to-side fluorescence ratios for GFP-tagged open-conformation SFK mutants. All mutants except Lyn show significant curvature enrichment compared to GFP–CaaX control. *n* = 18/13/17/12/15/12/14/13/16/15 cells (left-right), from two independent experiments. Data are mean ± standard deviation (SD); *p*-values from one-way ANOVA with Dunnett’s T3 multiple comparisons. **f**, Curvature localization of nine GFP-tagged human SFK mutants on nanobars. All except Lyn show enrichment at nanobar ends; all except Frk show localization to focal adhesions (see also Extended Data Fig. 1d). Scale bars, 10 µm. **g,** Gradient nanobar arrays were used to probe the curvature sensitivity of Src(pY419) enrichment. Top: bright-field image; middle: Src(pY419) immunostaining; bottom: averaged Src(pY419) signal per nanobar width across the array. Scale bar, 10 µm. **h,** Quantification shows that Src(pY419) curvature enrichment decreases as curvature becomes shallower, with significant enrichment at curvature diameters ≤ 500 nm. *n* = 8 arrays per condition (each containing 40 nanobars per width) from two independent experiments. Data are mean ± SD; *p*-values from one-sample t-test against ratio = 1.

To examine Src activity at curved membranes, we used an antibody recognizing Src phosphorylated at tyrosine 419 [Src(pY419)], a well-established marker of full kinase activation^55–58^. In this study, amino acid numbering refers to the human protein sequences unless otherwise indicated. On flat regions, Src(pY419) staining in A549 lung adenocarcinoma cells appeared faint and diffuse (**Fig. 1b**). In contrast, Src(pY419) was strongly enriched at nanopillars within the same field. Interestingly, in each cell, this enrichment occurred at only a subset of nanopillars, despite uniform membrane wrapping around all nanopillars as shown by the plasma-membrane marker GFP–CaaX (**Fig. 1b**). Similar curvature-dependent enrichment of Src(pY419) was observed across multiple cancer cell lines, U2OS (osteosarcoma), HeLa (cervical adenocarcinoma), MDA-MB-231 (breast adenocarcinoma), and BxPC3 (pancreatic adenocarcinoma) (**Extended Data Fig. 1a**).

To determine whether this enrichment reflects accumulation of total Src or specifically its active form, we expressed either wild-type Src–GFP or the open-conformation mutant Src(Y530F)–GFP. In this mutant, the autoinhibitory tyrosine (Y530) is replaced by phenylalanine, which cannot be phosphorylated, thereby favoring an open, activation-prone conformation. Cells were imaged 7 hours post-transfection to minimize detachment caused by prolonged expression of open-conformation Src^59,60^. Overexpressed wild-type Src–GFP, largely inactive under basal conditions, was evenly distributed across the plasma membrane without clear enrichment at curved regions (nanobar ends, arrowheads) (**Fig. 1c**). In contrast, Src(Y530F)–GFP showed strong enrichment at curved membranes (**Fig. 1d**). Consistent with previous studies^61,62^, Src(Y530F)–GFP also accumulated in focal adhesions at the cell periphery (arrows), confirmed by co-staining with vinculin, whereas vinculin was absent from curved membranes. These results indicate that membrane curvature drives preferential accumulation of active Src, independent of focal adhesion formation.

We next examined overall tyrosine phosphorylation using a pan-phosphotyrosine (pTyr) antibody. On flat regions, pTyr signals localized mainly to focal adhesions, consistent with their role as signaling hubs for focal adhesion kinase (FAK) and related kinases^63^. In contrast, on nanostructure regions, pTyr signals were observed both in focal adhesions and at curved membranes wrapping around nanopillars or nanobar ends (**Extended Data Fig. 1a**). These findings suggest that membrane curvature generates signaling domains that promote tyrosine kinase activation, functionally analogous but mechanistically distinct from focal adhesions. We refer to this mechanism as curvature-induced kinase activation (CIKA).

Since the Src(pY419) antibody recognizes a phosphorylation site within the activation loop that is highly conserved among SFKs (**Extended Data Fig. 1c**), we next examined whether curvature enrichment is a general property of active SFKs. We generated open-conformation mutants of all nine human SFKs, each tagged with C-terminal GFP: Src(Y530F), Yes(Y537F), Fyn(Y531F), Lck(Y505F), Hck(Y522F), Fgr(Y523F), Blk(Y501F), Frk(Y497F), and Lyn(Y508F). On flat regions, all mutants except Frk(Y497F) localized to focal adhesions (**Extended Data Fig. 1d**), consistent with previous reports of direct interactions between SFKs and FAK^64–66^. On nanobar arrays, all SFK mutants except Lyn(Y508F) were enriched at the curved nanobar ends (**Fig. 1e**,**f**), including Frk(Y497F), which did not localize to focal adhesions. These patterns suggest that distinct mechanisms govern the enrichment of active SFKs at curvature versus focal adhesions. Quantification of the fluorescence intensity ratio between nanobar ends and sides confirmed significant curvature enrichment (>1) for all open-conformation SFKs except Lyn. Among all SFKs, Src, Yes, and Fyn showed the strongest enrichment, whereas GFP–CaaX, used as a membrane control, displayed a ratio close to 1 (**Fig. 1e**).

We next asked whether the curvature enrichment of Src is related to curved adhesions, a recently identified adhesion structure that preferentially forms at curved membranes^34^. Src(Y530F) did not spatially correlate with integrin β5 (ITGB5), a marker of curved adhesions, on vitronectin-coated nanopillars (**Extended Data Fig. 1e**). Moreover, Src(Y530F) still accumulated at curved regions on gelatin-coated nanopillars. Gelatin is not a ligand for integrin β5 and does not support curved adhesion formation (**Extended Data Fig. 1f**). These findings indicate that Src enrichment at curved membranes occurs independently of curved adhesions.

To determine the curvature range that promotes Src enrichment, we used a gradient nanobar array with bar widths ranging from 100 to 1000 nm. Thinner bars generate higher curvature (1/diameter) at their ends (**Fig. 1g**). Src(pY419) strongly accumulated at the ends of the thinnest nanobars, with curvature enrichment decreasing progressively as bar width increased. Quantification of ∼300 nanobars per size across eight arrays confirmed this trend, revealing significant curvature-dependent Src activation at diameters ≤ 500 nm (**Fig. 1h**). This range differs from the broader curvature range (≤ 2500-nm diameter) reported for curved adhesion formation^34^.

### Membrane curvature-induced Src enrichment requires biomolecular condensation

TOCA-family proteins (FBP17, CIP4, and TOCA1) recognize high membrane curvature (≤ 500-nm diameter) through their conserved N-terminal F-BAR domains^67–69^. Their curvature range^67^ matches the range that enriches active Src. Since Src lacks an intrinsic curvature-sensing domain, we hypothesized that TOCA-family proteins recruit Src to curved membranes via their C-terminal SH3 domains, which are known to mediate protein–protein interactions (**Fig. 2a**)^70^.

**Fig. 2:**
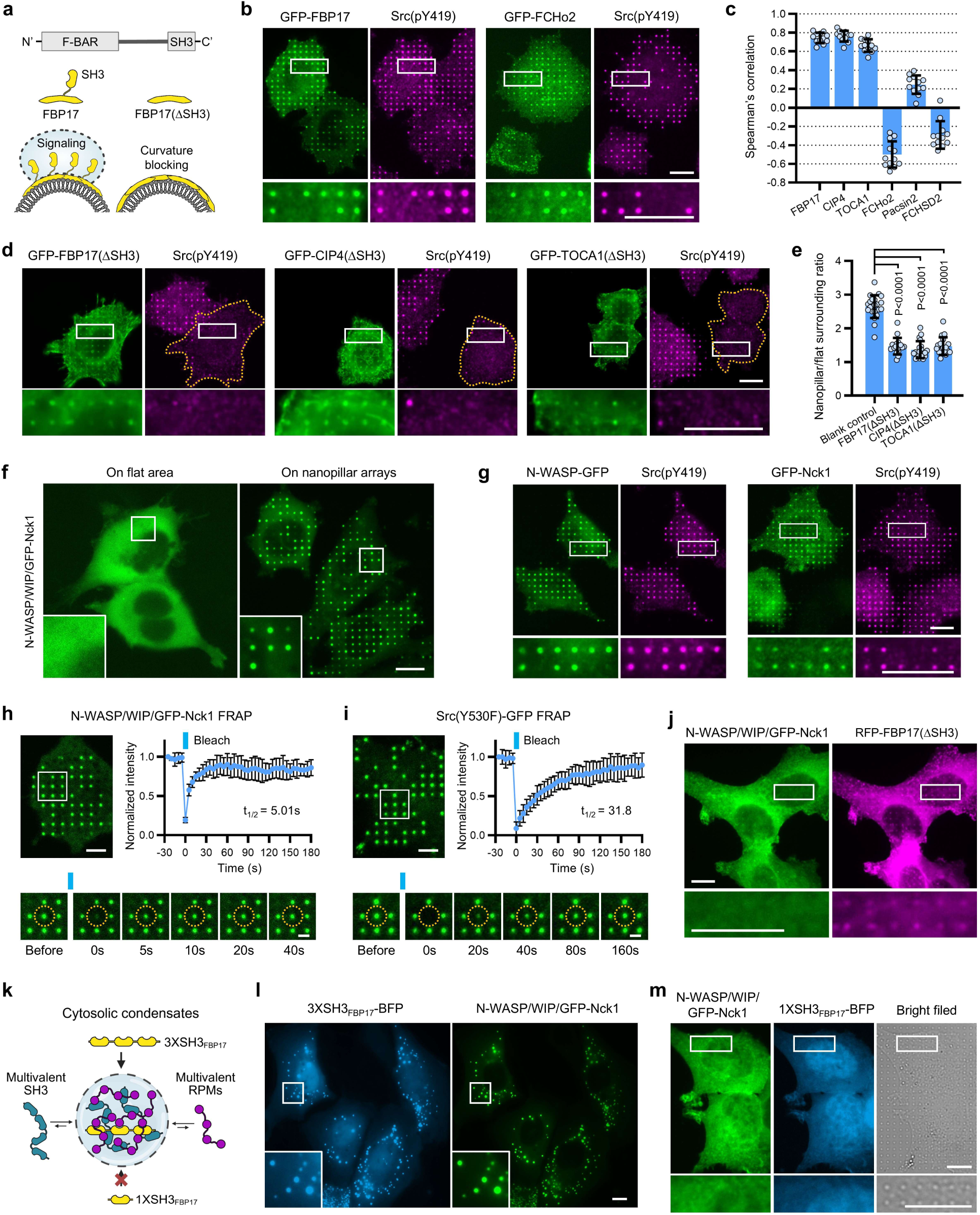
Membrane curvature-induced Src enrichment requires phase condensation. **a,** Domain organization of TOCA-family proteins, containing an N-terminal F-BAR domain for curvature sensing and a C-terminal SH3 domain for protein interactions. SH3-deletion mutants, such as FBP17(ΔSH3), lack the interaction module required for downstream signaling. **b,** GFP–FBP17 co-accumulates with active Src (Src(pY419)) at nanopillars, while GFP-FCHo2 shows no correlation or even anti-correlation with Src activation at nanopillars. Scale bars: 10 µm. **c,** Spearman’s correlation coefficients between various F-BAR proteins and Src(pY419) at nanopillars. TOCA-family proteins (FBP17, CIP4, TOCA1) correlate strongly with Src(pY419), whereas other F-BAR proteins (FCHo2, Pacsin2, FCHSD2) do not. *n* = 12 cells per condition, from two independent experiments. **d,** Expression of SH3-deletion mutants (FBP17(ΔSH3), CIP4(ΔSH3), TOCA1(ΔSH3)) abolishes Src(pY419) enrichment at nanopillars. Scale bars: 10 µm. **e,** Quantification confirms significant reduction of Src(pY419) enrichment at nanopillars with the expression of SH3-deletion mutants. *n* = 22, 17, 20, and 18 cells; two independent experiments. Data are mean ± SD; *p*-values from Kruskal-Wallis test with Dunn’s multiple comparisons. **f,** GFP–Nck1, co-expressed with N-WASP and WIP, is diffuse on flat membranes but is concentrated at nanopillars. Scale bar: 10 µm. **g,** GFP–Nck1 and N-WASP–GFP both co-accumulate with Src(pY419) at nanopillars, indicating the incorporation of Src(pY419) into curvature-induced condensates. Scale bars: 10 µm. **h** and **i,** FRAP analyses show rapid fluorescence recovery for GFP–Nck1 (t_1/2_ ∼5 s) and for Src(Y530F)–GFP (t_1/2_ ∼32 s), indicating liquid-like dynamics of curvature-induced condensates. Scale bars: 10 µm; 2 µm (montage panels). *n* = 5 nanopillars from 5 independent cells per condition. See also Supplementary Video 1,2. **j**, Expression of FBP17(ΔSH3) abolishes the formation of N-WASP–WIP/GFP–Nck1 condensates at nanopillars Scale bars, 10 µm. **k,** Schematic of a trivalent SH3 construct (3×SH3_FBP17_–BFP) designed to induce phase condensation through multivalent SH3–PRM interactions, and a monovalent SH3 construct (1×SH3_FBP17_–BFP) designed to disrupt phase condensation by competitively inhibiting the multivalent interactions. **l,** 3×SH3_FBP17_–BFP co-condenses with N-WASP/WIP/GFP–Nck1, forming cytosolic droplets. Scale bar, 10 µm. **m**, A monovalent SH3 construct (1×SH3_FBP17_–BFP) disrupts the formation of N-WASP/WIP/GFP–Nck1 condensates at nanopillars. Scale bars, 10 µm.

Supporting this hypothesis, Src(pY419) was strongly enriched at nanopillars where GFP–FBP17 accumulated (**Fig. 2b**), with Spearman’s correlation analysis confirming this high spatial correlation (**Fig. 2c**). CIP4 and TOCA1 showed similar curvature accumulation and spatial correlation with Src(pY419) (**Extended Data Fig. 2a** and **Fig. 2c**). In contrast, curvature-sensing proteins from other F-BAR families, such as FCHo2 (**Fig. 2b**), Pacsin2, and FCHSD2 (**Extended Data Fig. 2b**), showed little to no correlation or even anti-correlation with Src(pY419) (quantification in **Fig. 2c**), suggesting that curvature-induced Src enrichment is specific to the TOCA family.

The selective enrichment of pSrc at certain nanopillars, but not others, indicate a digital rather than graded response. Such behavior is consistent with a condensation mechanism. TOCA family proteins contain an SH3 domain, which is known to scaffold biomolecular condensation through multivalent interactions with proline-rich motifs^53^. Although each TOCA-family protein contains only a single SH3 domain, they cluster and oligomerize at curved membranes^68,69^, thereby increasing local SH3 valency. We hypothesized that curvature-driven TOCA clustering promotes condensate formation via multivalent SH3–PRM interactions, which in turn recruits Src.

To test the contribution of TOCA-family SH3 domains to Src enrichment, we generated SH3-deletion mutants FBP17(ΔSH3), CIP4(ΔSH3), and TOCA1(ΔSH3), which retain curvature binding but lack SH3-mediated interactions (**Fig. 2a**). Expression of each ΔSH3 mutant nearly abolished Src(pY419) enrichment at nanopillars (**Fig. 2d,e**). These ΔSH3 mutants also specifically reduced pTyr accumulation at curved membranes without affecting pTyr signals in focal adhesions (**Extended Data Fig. 2c**). These results indicate that the SH3 domains are essential for curvature-induced kinase activation.

To examine the SH3-mediated condensation hypothesis, we focused on the Nck–N-WASP–WIP complex^71^. The Nck and N-WASP forms a multivalent SH3–PRM network that is known to condensate *in vitro*^53^, and the N-WASP–WIP complex has been reported to interact with TOCA-family proteins^72,73^. On flat regions, GFP–Nck1 appeared diffusive in the cytosol of cells co-expressing GFP–Nck1, N-WASP, and WIP, consistent with previous reports that cellular concentrations of these proteins are below the condensation threshold^53^. In contrast, GFP–Nck1 showed all-or-none enrichment at certain nanopillar locations (**Fig. 2f**). Importantly, both N-WASP–GFP and GFP–Nck1 were enriched on these nanopillars where Src(pY419) accumulated (**Fig. 2g**). This was further supported by the curvature enrichment of endogenous Nck1 and N-WASP in untransfected cells, which also colocalized with Src(pY419) (**Extended Data Fig. 2d**). These results suggest that membrane curvature can drive Nck–N-WASP–WIP condensation at endogenous concentrations and that active Src is incorporated into these condensates.

Fluorescence recovery after photobleaching (FRAP) revealed rapid molecular exchange between the curvature-induced condensates and the surrounding cytosol, with half-times of ∼5 s for GFP–Nck1 (**Fig. 2h** and **Supplementary Video 1**) and ∼32 s for Src(Y530F)–GFP (**Fig. 2i** and **Supplementary Video 2**). These dynamics indicate that these condensates exhibit liquid-like properties.

As expected, expression of FBP17(ΔSH3) abolished GFP–Nck1 accumulation at nanopillars in cells co-expressing N-WASP and WIP (**Fig. 2j**), confirming that SH3 domains of TOCA-family proteins are required for curvature-induced condensation.

We next asked whether SH3 clustering alone can drive Nck–N-WASP–WIP condensation. To test this, we designed a trivalent construct (3xSH3_FBP17_–BFP) containing three tandem SH3 domains from FBP17 fused to a blue fluorescence protein (BFP) (**Fig. 2k**). When coexpressed with GFP–Nck1, N-WASP, and WIP, this construct induced robust cytosolic condensate formation (**Fig. 2l**), producing dynamic droplets that exhibited rapid FRAP (**Extended Data Fig. 2e** and **Supplementary Video 3**). These droplets also underwent frequent fusion and fission, confirming their liquid-like properties. In contrast, a similar trivalent construct containing three SH3 domains from Nck1 (3xSH3_Nck1_–BFP) failed to induce condensation under identical conditions (**Extended Data Fig. 2f**), indicating that clustering of FBP17’s SH3 domains, but not Nck1’s SH3 domains, can drive Nck–N-WASP–WIP condensation in cells.

To confirm the role of multivalent interactions in condensation, we generated a monovalent SH3 construct, 1xSH3_FBP17_–BFP, which competitively blocks SH3–PRM binding (**Fig. 2k**). When coexpressed with GFP–Nck1, N-WASP, and WIP, this construct was diffuse (**Extended Data Fig. 2g**), and strongly suppressed curvature-induced condensate formation at nanopillars (**Fig. 2m**). It also markedly reduced pTyr at curved membranes, without affecting pTyr in focal adhesions (**Extended Data Fig. 2h**). Although the inhibitory effect of 1xSH3_FBP17_ resembled that of FBP17(ΔSH3) (**Fig. 2j** and **Extended Data Fig. 2c**), the two act through distinct mechanisms: FBP17(ΔSH3) acts as a curvature binder lacking SH3-mediated scaffolding capacity, whereas 1xSH3_FBP17_ competitively disrupts the multivalent interactions required for condensation. Together, these findings support a model in which membrane curvature promotes phase condensation by clustering SH3-containing TOCA family proteins, a process necessary for recruiting and activating Src at curved membranes.

### Curvature-induced condensates recruit Src via its SH3–SH2 domains and adaptor proteins

Src contains four domains, an N-terminal SH4 domain for membrane anchoring, central SH3 and SH2 domains for protein interactions, and a C-terminal kinase domain (SH1)^44,45^. To determine which domains govern curvature enrichment, we generated a series of Src(Y530F) truncation mutants lacking each domain (**Fig. 3a**). Src(ΔSH4, Y530F) was largely cytosolic; however, following fixation and permeabilization to reduce the diffusive background, Src(ΔSH4, Y530F) showed accumulation at focal adhesions and nanobar ends (**Fig. 3b**,**c**), indicating that membrane anchoring is not essential for either localization. By contrast, Src(ΔSH3, Y530F) retained focal adhesion localization but completely lost enrichment at nanobar ends, demonstrating that the SH3 domain is specifically required for curvature targeting. Src(ΔSH2, Y530F) failed to localize to either site, indicating that the SH2 domain is required for both curvature and focal adhesion localization. Finally, Src(ΔSH1) maintained strong enrichment at both focal adhesions and curved membranes, showing that the kinase domain is dispensable for either localization.

**Fig. 3:**
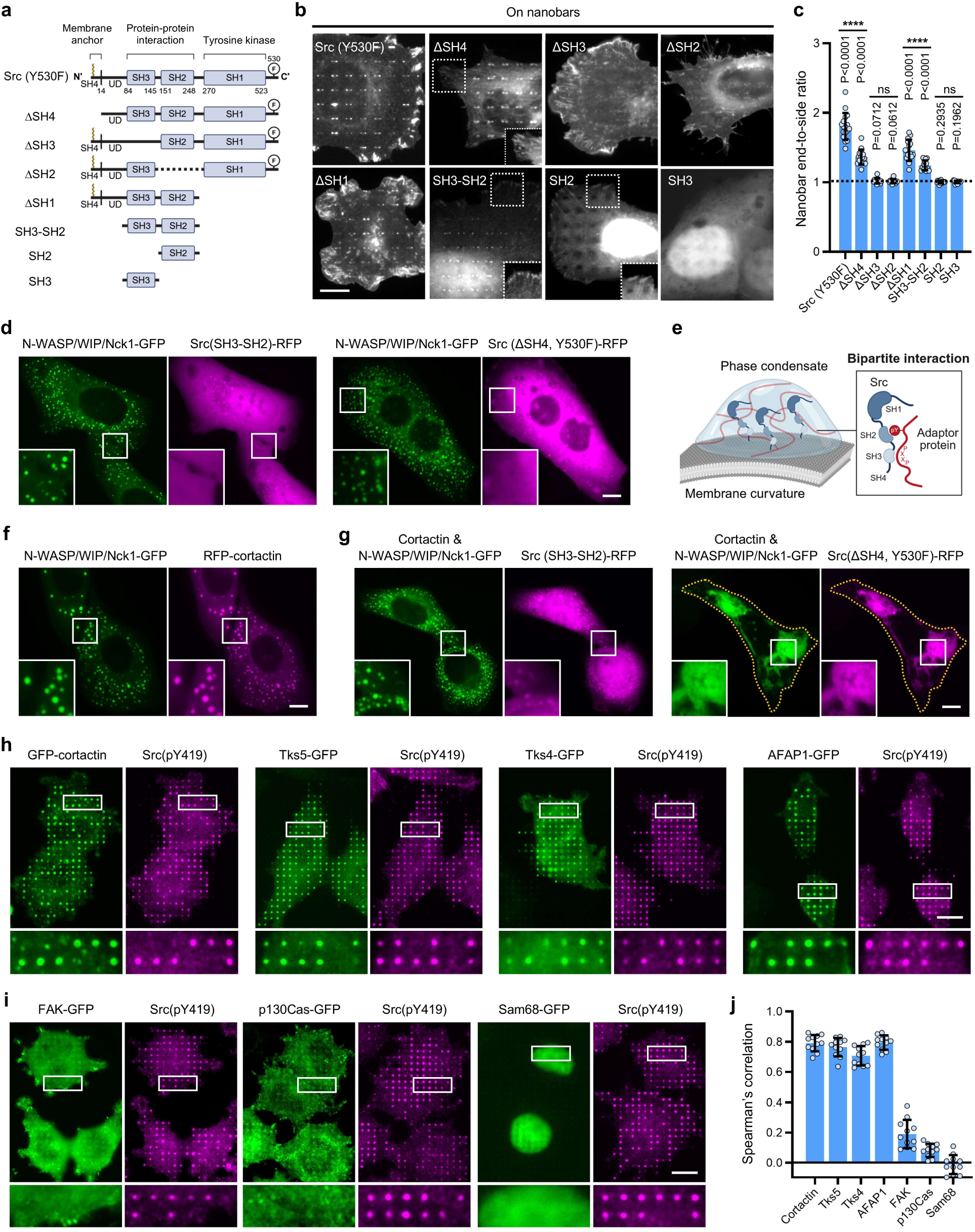
Curvature-induced condensates recruit Src via its SH3–SH2 domains and adaptor proteins. **a,** Domain structure of Src(Y530F) showing SH4 (membrane anchor), SH3, SH2, and SH1 (kinase) domains. Schematic shows the design of truncation constructs. **b,** Representative images of the truncation constructs expressed in cells cultured on nanobar arrays. Curvature recruitment requires both SH3 and SH2 domains, whereas focal adhesion localization depended solely on the SH2 domain. The intensity in the dotted boxes is increased to highlight focal adhesions. Scale bar, 10 µm. **c,** Quantification of curvature enrichment (nanobar end-to-side fluorescence ratio) for constructs in (B). *n* = 17, 15, 12, 13, 15, 14, 16, and 15 cells from two independent experiments; data are mean ± SD; *p*-values from one-sample t-test against ratio = 1. **d,** Src(SH3–SH2) and Src(ΔSH4, Y530F) are not incorporated into cytosolic condensates formed by 3×SH3_FBP17_–BFP, Nck1–GFP, N-WASP, and WIP. See Extended Data Fig. 3a for the BFP channel. Scale bar, 10 µm. **e,** Schematic illustrating how adaptor proteins containing both PRMs and phosphotyrosine motifs engage the SH3 and SH2 domains of Src in a bipartite manner, thereby recruiting Src into condensates. **f**, Cortactin is efficiently incorporated into cytosolic condensates assembled by 3×SH3_FBP17_–BFP, GFP–Nck1, N-WASP, and WIP. See Extended Data Fig. 3b for the BFP channel. Scale bar, 10 µm. **g,** Co-expression of cortactin with 3×SH3_FBP17_–BFP/N-WASP/WIP/GFP–Nck1 enables incorporation of Src(SH3–SH2) and Src(ΔSH4, Y530F) into condensates. Cortactin with Src(ΔSH4, Y530F) produces larger condensates. See Extended Data Fig. 3c for the BFP channel. Scale bar, 10 µm. **h,** Cortactin, Tks4, Tks5, and AFAP1 colocalize with Src(pY419) at nanopillars, supporting their role as curvature-enriched adaptor proteins. Scale bar: 10 µm. **i**, FAK and p130Cas localize to focal adhesions, and Sam68 is nuclear; none show enrichment at nanopillars. Scale bar, 10 µm. **j,** Spearman’s correlation coefficients between adaptor proteins and Src(pY419) at nanopillars. Cortactin, Tks4, Tks5, and AFAP1 show strong correlations with curvature-enriched Src, whereas FAK, Sam68, and p130Cas do not. *n* = 12 cells per condition, from two independent experiments; data are mean ± SD.

To further demonstrate domain contributions, we expressed the SH3–SH2 tandem, SH3, and SH2 constructs of Src (**Fig. 3a**). After fixation and permeabilization, fluorescence imaging revealed that SH3-SH2 localized to both focal adhesions and curved membranes; SH2 alone targeted only focal adhesions; SH3 alone showed no enrichment at either site (**Fig. 3b**,**c**). These results indicate that the SH2 domain is sufficient for focal adhesion targeting, whereas curvature recruitment requires cooperative input from both SH3 and SH2 domains.

We next tested whether Src(SH3–SH2) could be incorporated into reconstituted cytosolic condensates formed by 3×SH3_FBP17_–BFP, GFP–Nck1, N-WASP, and WIP. Surprisingly, Src(SH3–SH2) did not accumulate in these condensates in live cells (**Fig. 3d** and **Extended Data Fig. 3a**). Likewise, the cytosolic mutant Src(ΔSH4, Y530F)-RFP showed no detectable enrichment in these condensates. These results indicate that the SH3–SH2 domains are necessary but not sufficient for their condensate recruitment.

Src interacts with multivalent adaptor proteins such as cortactin, which contains both PRMs and phosphotyrosine sites that can simultaneously engage Src SH3 and SH2 domains in a bipartite manner (**Fig. 3e**)^66,74–79^. To test whether such adaptors mediate Src recruitment into condensates, we co-expressed cortactin together with 3×SH3_FBP17_, GFP–Nck1, N-WASP, and WIP. Cortactin was strongly incorporated into the condensates (**Fig. 3f** and **Extended Data Fig. 3b**). When Src(SH3–SH2) was co-expressed with cortactin and the condensation machinery, it showed weak but detectable recruitment into condensates (**Fig. 3g** and **Extended Data Fig. 3c**). Surprisingly, co-expression of Src(ΔSH4, Y530F), which retains the kinase domain, together with the same condensation components, triggered dramatic condensate growth, producing unusually large structures (**Fig. 3g** and **Extended Data Fig. 3c**). This observation suggests that Src kinase activity amplifies condensate formation, likely by phosphorylating phase components and creating additional interaction sites. For example, Src-mediated cortactin phosphorylation generates binding sites for the SH2 domain of Nck^80^, which could promote condensate growth. These enlarged condensates remained liquid-like, as shown by the rapid FRAP recovery of Nck1 and Src(ΔSH4, Y530F) (**Extended Data Fig. 3d** and **Supplementary Video 4**).

To identify adaptor proteins that contribute to Src enrichment at curvature, we examined seven candidates previously reported to interact with Src through bipartite engagement of its SH3 and SH2 domains: cortactin, Tks4, Tks5, AFAP1, FAK, Sam68, and p130Cas (**Extended Data Fig. 3e**)^66,74–79^. Among these, cortactin, Tks4, Tks5, and AFAP1 showed strong curvature preference at nanobar ends, whereas FAK, p130Cas and Sam68 showed no curvature enrichment (**Extended Data Fig. 3f**-**h**). Consistent with these results, Tks4, Tks5, and AFAP1 were strongly enriched at nanopillars and spatially correlated with Src(pY419) (**Fig. 3h**). Notably, Src(pY419) colocalized with cortactin and AFAP1 specifically at nanopillars, but not with cortactin puncta between nanopillars or AFAP1 associated with stress fibers (**Fig. 3h**). In contrast, FAK and p130Cas were confined to focal adhesions and absent from nanopillars, whereas Sam68 was predominantly nuclear; none correlated with Src(pY419) at curved membranes (**Fig. 3i**). Spearman’s correlation analysis confirmed that, at curved membranes, active Src strongly correlates with cortactin, Tks4, Tks5, and AFAP1, but not with FAK, p130Cas, or Sam68 (**Fig. 3j**).

Together, these results indicate that curvature-induced condensates recruit Src through bipartite engagement of its SH3 and SH2 domains with a subset of multivalent adaptor proteins (cortactin, Tks4, Tks5, and AFAP1).

### Curvature-induced condensation promotes local Src activation and downstream PI3K/Akt signaling

Since Src autophosphorylation at Y419 is intermolecular and concentration-dependent^81,82^, we hypothesized that membrane curvature not only enriches Src in its open conformation but also promotes its local activation by facilitating Y419 phosphorylation (**Fig. 4a**). To distinguish between selective enrichment and activation, we immunostained for Src(pY419) in cells expressing either wild-type Src-GFP or the open-conformation mutant Src(Y530F)–GFP. Wild-type Src-GFP exhibited diffuse localization with modest enrichment at nanopillars, whereas Src(pY419) almost exclusively localized at nanopillars in the same cells (**Fig. 4b**). Src(Y530F)-GFP localized to both nanopillars and focal adhesions, but the corresponding Src(pY419) signal was markedly higher at nanopillars. Pixel-by-pixel heatmaps of the activation ratios [Src(pY419)/Src and Src(pY419)/Src(Y530F), normalized to flat membrane levels] confirmed significantly elevated Src activation at nanopillars (**Fig. 4b**,**c**). Although Src(Y530F) strongly accumulated at focal adhesions, its activation ratio there was only modestly elevated (∼1.3), whereas the ratio at nanopillars was substantially higher (∼2.0). These results support that membrane curvature promotes local Src activation beyond mere enrichment.

**Fig. 4:**
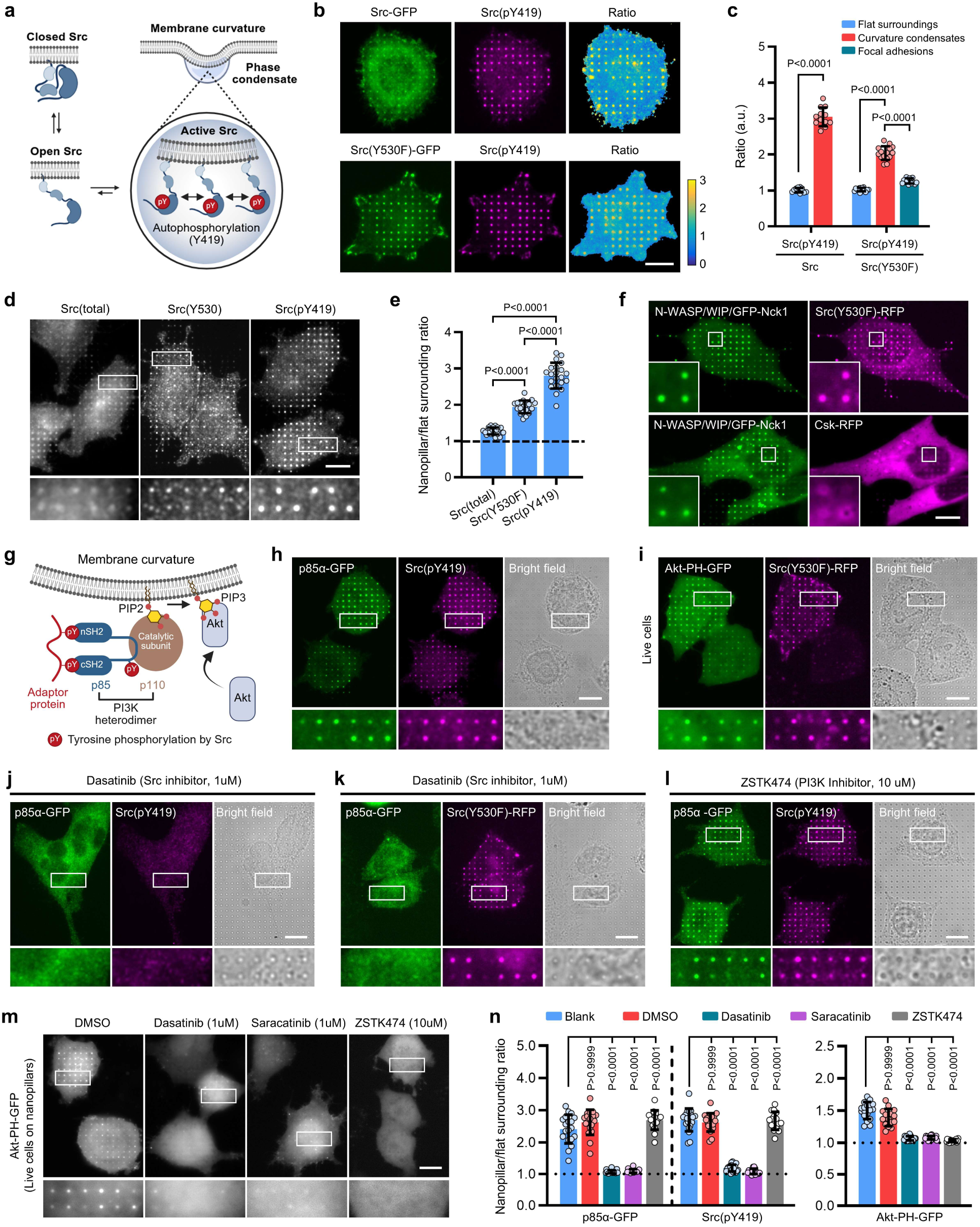
Curvature-induced condensation promotes local Src activation and PI3K/Akt signaling. **a,** Schematic illustrating curvature-induced Src activation: condensates formed at membrane curvature enrich open-conformation Src, promoting local trans-autophosphorylation at Y419. **b,** Ratiometric imaging of Src(pY419)/Src–GFP and Src(pY419)/Src(Y530F)–GFP reveals enhanced Y419 phosphorylation at nanopillars. Scale bar, 10 µm. **c,** Quantification confirms that Src autophosphorylation (pY419) is significantly elevated at nanopillars compared to flat regions or focal adhesions. *n* = 12 (Src–GFP) and 15 (Src(Y530F)–GFP) cells, from two independent experiments; data are mean ± SD; *p* values from one-way ANOVA with Dunnett’s T3 multiple comparisons. **d,** Immunostaining for total Src, open conformation Src(Y530), and active Src(pY419) shows different degrees of enrichment at nanopillars. Scale bar, 10 µm. **e,** Quantification reveals highest curvature enrichment for Src(pY419), intermediate for Src(Y530), and minimal for total Src. *n* = 22, 19, 21 cells from two independent experiments; mean ± SD; *p* values from one-way ANOVA with Dunnett’s T3 multiple comparisons. **f,** At nanopillars, Src(Y530F)–RFP colocalizes with N-WASP/WIP/GFP–Nck1 condensates, while Csk–RFP (a negative regulator of Src) is excluded. Scale bar, 10 µm. **g,** Schematic illustrating PI3K/Akt signaling downstream of curvature-induced Src activation. **h,** PI3K regulatory subunit alpha (p85α–GFP) accumulates at nanopillars, colocalizing with Src(pY419). Scale bar, 10 µm. **i,** Live-cell imaging showing Akt–PH–GFP, a PI(3,4,5)P3 sensor, enriched at nanopillars together with Src(Y530F)–RFP. Scale bar, 10 µm. **j,** Treatment with Src inhibitor dasatinib abolishes p85α–GFP and Src(pY419) enrichment at nanopillars. Scale bar, 10 µm. **k,** Dasatinib treatment eliminates p85α–GFP enrichment at nanopillars, while Src(Y530F)–RFP enrichment remains, indicating that curvature localization of Src(Y530F) does not rely on its kinase activity. Scale bar, 10 µm. **l,** PI3K inhibition (ZSTK474) does not affect Src(pY419) or p85α–GFP enrichment at nanopillars. Scale bar, 10 µm. **m,** Enrichment of Akt–PH–GFP at nanopillars is abolished by either Src or PI3K inhibition, indicating that both kinases are required for Akt signaling at curvature. Scale bar, 10 µm. **n,** Quantification of curvature enrichment for p85α-GFP, Src(pY419), and Akt–PH–GFP under the indicated inhibitor treatments. *n* = 19, 18, 21, 17, and 16 cells (p85α-GFP co-stained with Src(pY419)); 20, 15, 17, 19, and 19 cells (Akt–PH–GFP), from two independent experiments; data are mean ± SD; *p*-values from Kruskal-Wallis tests with Dunn’s multiple comparisons.

To confirm that membrane curvature enhances Src activation at endogenous expression levels, we examined Src distribution and activation using three antibodies recognizing total Src, open-conformation Src (Src(Y530)), and fully active Src (Src(pY419)). All three signals were enriched at nanopillars, but to varying extents (**Fig. 4d**). Quantification showed the strongest enrichment for Src(pY419), followed by Src(Y530), and the weakest for total Src (**Fig. 4e**), supporting that curvature enhances local Src activation under endogenous conditions.

To explore additional mechanisms that facilitate local Src activation, we examined the spatial distribution of Csk, a key negative regulator that phosphorylates Src at Y530 and promotes its autoinhibition^83^. While Src(Y530F)–RFP accumulated and colocalized with Nck1-marked condensates at nanopillars, Csk–RFP was selectively excluded from these condensates (**Fig. 4f**). Thus, curvature-induced condensates not only concentrate Src for autophosphorylation but also spatially exclude its negative regulator, creating a microenvironment that favors Src activation.

We next asked whether Src activation at curvature propagates to downstream PI3K/Akt signaling, a key oncogenic pathway. Src is known to recruit PI3K, a heterodimer composed of a regulatory p85 and a catalytic p110 subunit, to the plasma membrane by generating phosphotyrosine docking sites for the nSH2–cSH2 domains of p85 (Ref. ^84^). In addition, Src can also directly phosphorylate p85 to relieve inhibition of p110 (Fig. 4g)^85,86^.

Supporting this model, p85ɑ–GFP strongly accumulated at nanopillars and spatially correlated with Src(pY419) (**Fig. 4h**). The nSH2–cSH2 fragment alone was sufficient for curvature recruitment and correlation with Src(pY419) (**Extended Data Fig. 4a**). Furthermore, nanopillar-localized p85α was phosphorylated at Y467 (p85α(pY467); human numbering) (**Extended Data Fig. 4b**), a modification known to increase PI3K activity^86^. As a functional readout of PI3K, we used a GFP-tagged Akt PH domain (Akt-PH-GFP), which binds PI(3,4,5)P3, the lipid product of active PI3K. In live cells, Akt-PH-GFP accumulated at nanopillars in correlation with Src(Y530F)-RFP (**Fig. 4i**).

To confirm that curvature-induced PI3K/Akt signaling requires Src kinase activity, we treated cells with the Src inhibitors dasatinib or saracatinib (1 µM). Both inhibitors abolished nanopillar enrichment of p85α–GFP and Src(pY419) (**Fig. 4j**, **Extended Data Fig. 4c** and quantification in **Fig. 4n**). However, neither inhibitor disrupted nanopillar localization of Src(Y530F)-RFP, even though they abolished Src(pY419) in the same cells (**Fig. 4k** and **Extended Data Fig. 4d**). Moreover, both inhibitors eliminated the p85α(pY467) signal at nanopillars (**Extended Data Fig. 4e**). These results indicate that Src kinase activity is dispensable for its own recruitment to curvature but essential for activating PI3K, thereby establishing Src as the upstream regulator of PI3K in curvature-dependent signaling.

Conversely, inhibition of PI3K with ZSTK474 (10 µM) had no effect on the curvature enrichment of Src(pY419) or p85α-GFP at nanopillars (**Fig. 4l** and quantification in **4n**), confirming that PI3K activity is not required for Src activation or for p85α recruitment at curved membranes. By contrast, Akt-PH–GFP enrichment at nanopillars was abolished by either Src or PI3K inhibition (**Fig. 4m** and quantification in **4n**), indicating that curvature-dependent PI(3,4,5)P3 production requires both upstream Src and PI3K activity. Taken together, these results demonstrate that membrane curvature promotes localized Src activation, which in turn, recruits and activates PI3K and Akt at curved membranes.

### CIKA promotes anchorage-independent cell survival

Most mammalian cells are anchorage-dependent, relying on cell adhesions to activate pro-survival pathways such as FAK–Src, PI3K/Akt, and MAPK/ERK. Loss of adhesion triggers anoikis, preventing detached cells from surviving^12,13^. During metastasis, tumor cells often acquire resistance to anoikis, enabling anchorage-independent survival, a hallmark of metastatic potential. One striking feature of metastatic cells is their extensive plasma membrane folding and curvature, often localized to actin-rich regions^23,27^. Given that Src is an important player in anoikis resistance^87^, we hypothesized that regions of high membrane curvature in suspended cancer cells serve as alternative signaling hubs that promote Src activation and survival in the absence of adhesion.

To test this hypothesis, we examined Src activation (Src(pY419)) and actin organization in A549 cells under three conditions: adhered on flat substrates (**Fig. 5a**), adhered on nanopillar substrates (**Fig. 5b**), and maintained in suspension (**Fig. 5c**). On flat substrates, cells exhibited strong actin stress fibers but minimal Src(pY419). On nanopillar substrates, actin accumulated around nanopillars and spatially correlated with Src(pY419). In suspension, 3D confocal imaging revealed that cortical actin often polarized into crown-like structures (**Fig. 5c**), which contain pronounced membrane protrusions and invaginations^27^. Src(pY419) was not uniformly distributed along the cell cortex (**Extended Data Fig. 5a**). Instead, it was specifically enriched in curvature-rich crown regions (**Fig. 5c**), localizing preferentially at the bases of the crowns, sites of inward positive membrane curvature, rather than at the crown tips (**Fig. 5d**).

**Fig. 5:**
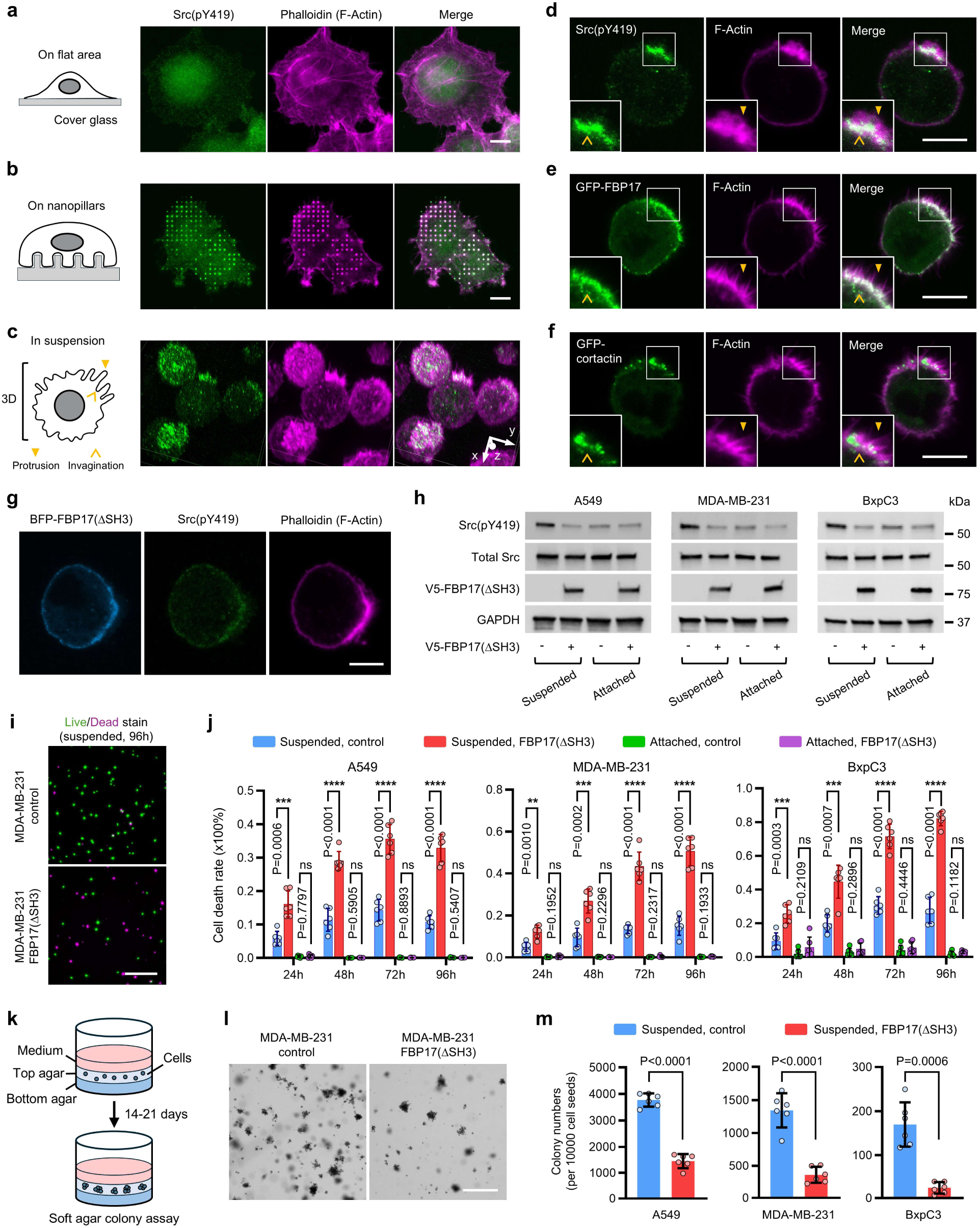
CIKA promotes anchorage-independent cell survival. **a,** On flat substrates, A549 cells exhibit prominent actin stress fibers (phalloidin) but minimal Src activation, as indicated by Src(pY419) staining. Scale bar, 10 µm. **b**, On nanopillar substrates, both actin and Src(pY419) strongly co-accumulate at sites of membrane curvature. Scale bar, 10 µm. **c,** 3D confocal reconstruction of suspended A549 cells reveals polarized cortical actin forming crown-like structures, where Src(pY419) is selectively enriched. Scale bars, 10 µm. **d-f,** 2D confocal slices of individual suspended cells showing that Src(pY419) (d), GFP–FBP17 (e), and GFP–cortactin (f) localize predominantly at crown-base invaginations rather than protrusive tips. Scale bars, 10 µm. **g,** Expression of BFP–FBP17(ΔSH3) abolishes Src(pY419) enrichment in suspended cells, confirming that CIKA is required for curvature-induced Src activation. Scale bar, 10 µm. **h,** Western blot analysis showing elevated Src(pY419) levels, despite similar total Src levels, in suspended versus adherent A549, MDA-MB-231, and BxPC3 cells. Expression of FBP17(ΔSH3) suppresses Src(pY419) in suspension but has little effect under adherent conditions. **i,** Live/dead staining showing that FBP17(ΔSH3) expression reduces survival of MDA-MB-231 cells in suspension (soft agar) but does not affect survival under adherent conditions. See also Extended Data Fig. 5e for adherent MDA-MB-231 cells as well as A549 and BxPC3 cells under both conditions. Scale bar, 500 µm. **j,** Quantification of cell viability showing that FBP17(ΔSH3) expression increases cell death in suspension but not under adherent conditions, across A549, MDA-MB-231, and BxPC3 cells. *n* = 6 independent cultures per condition, from two independent experiments; mean ± SD; *p*-values from t tests with Welch correction (parametric) and Mann Whitney tests (non-parametric). **k,** Schematic of the soft agar colony formation assay used to assess anchorage-independent colony formation. **l,** Representative images of MDA-MB-231 colony formation in soft agar. FBP17(ΔSH3)-expressing cells formed markedly fewer colonies than controls. Scale bars, 500 µm. See also Extended Data Fig. 5f for A549 and BxPC3 colony formation. **m,** Quantification of colony numbers (size threshold, 100-µm Feret diameter) showing that FBP17(ΔSH3) significantly reduces soft agar colony formation in all three cell lines. *n* = 6 independent cultures per condition, from two independent experiments; mean ± SD; *p*-values from t tests with Welch correction.

The curvature-sensing proteins FBP17 and CIP4 also accumulated at crown bases (**Fig. 5e** and **Extended Data Fig. 5b**). Consistently, the multivalent adaptor proteins cortactin and Tks5 were selectively enriched at crown bases (**Fig. 5f** and **Extended Data Fig. 5c**). In non-polarized suspended cells lacking a dominant crown, Src(pY419), FBP17, CIP4, and cortactin were likewise enriched at scattered invaginations across the cell cortex (**Extended Data Fig. 5d**). Expression of the dominant-negative construct FBP17(ΔSH3) markedly reduced Src(pY419) accumulation in suspended cells (**Fig. 5g**). Together, membrane invaginations in suspended cancer cells likely trigger signaling cascades similar to those elicited by nanopillar-induced membrane invaginations.

Western blotting revealed significantly higher levels of Src(pY419) in suspended cells compared with adherent cells, despite similar total Src levels (**Fig. 5h**). A similar increase in Src activation upon detachment has been reported in multiple metastatic cell types^14–16,18^. Notably, expression of FBP17(ΔSH3) markedly suppressed Src activation in suspended cells but had little to no effect under adherent conditions. These effects were consistent across A549, MDA-MB-231, and BxPC3 cell lines (**Fig. 5h**), suggesting CIKA as a general mechanism of Src activation in suspended cancer cells.

To assess the role of CIKA in cancer cell survival, we stably expressed BFP–FBP17(ΔSH3) or control BFP in A549, MDA-MB-231, and BxPC3 cells. Viability was measured under both adherent and suspension conditions at 24, 48, 72, and 96 hours. To prevent cell aggregation or reattachment, suspended cells were embedded in 0.35% soft agar. Live/dead staining showed that blocking CIKA by FBP17(ΔSH3) significantly reduced the survival of suspended, but not adherent, cells across all three cell lines (**Fig. 5i** and **Extended Data Fig. 5e**). Quantification revealed a progressive, time-dependent cell death induced by suspension and FBP17(ΔSH3) expression increased it (**Fig. 5j**). After 96 hours in suspension, cells expressing FBP17(ΔSH3) showed ∼82% death in BxPC3, ∼51% in MDA-MB-231, and ∼33% in A549. In contrast, survival of adherent cells was unaffected by FBP17(ΔSH3) expression at any time point, indicating that CIKA is important for anchorage-independent, but not anchorage-dependent, survival of cancer cells.

We further evaluated the role of CIKA using a soft-agar colony formation assay (**Fig. 5k**), an established *in vitro* model for assessing tumorigenic potential^88^. Equal numbers of single cells were seeded in agar and cultured in suspension for 14 days (A549, MDA-MB-231) or 21 days (BxPC3). Expression of FBP17(ΔSH3) markedly reduced colony formation in all three cell lines (**Fig. 5l** and **Extended Data Fig. 5f**). After applying a size threshold to exclude single cells and debris, quantification revealed that FBP17(ΔSH3) reduced colony numbers by ∼86% in BxPC3, ∼73% in MDA-MB-231, and ∼61% in A549 compared with control cells (**Fig. 5m**). These results demonstrate that CIKA inhibition suppresses anchorage-independent growth.

### Blocking CIKA suppresses cancer metastasis *in vivo*

To assess the role of CIKA in cancer metastasis *in vivo*, we used two established xenograft models, in which NSG mice were injected with human cancer cells either intraperitoneally to model peritoneal colonization^89,90^, or via the tail vein to model hematogenous lung metastasis^91,92^. Cancer cells were engineered to stably express either Antares (NanoLuc luciferase fused to the cyan-excitable orange fluorescent protein CyOFP1)^93^ or firefly luciferase for bioluminescent tracking, together with either FBP17(ΔSH3) to block CIKA or an empty control vector (**Fig. 6a**).

**Fig. 6:**
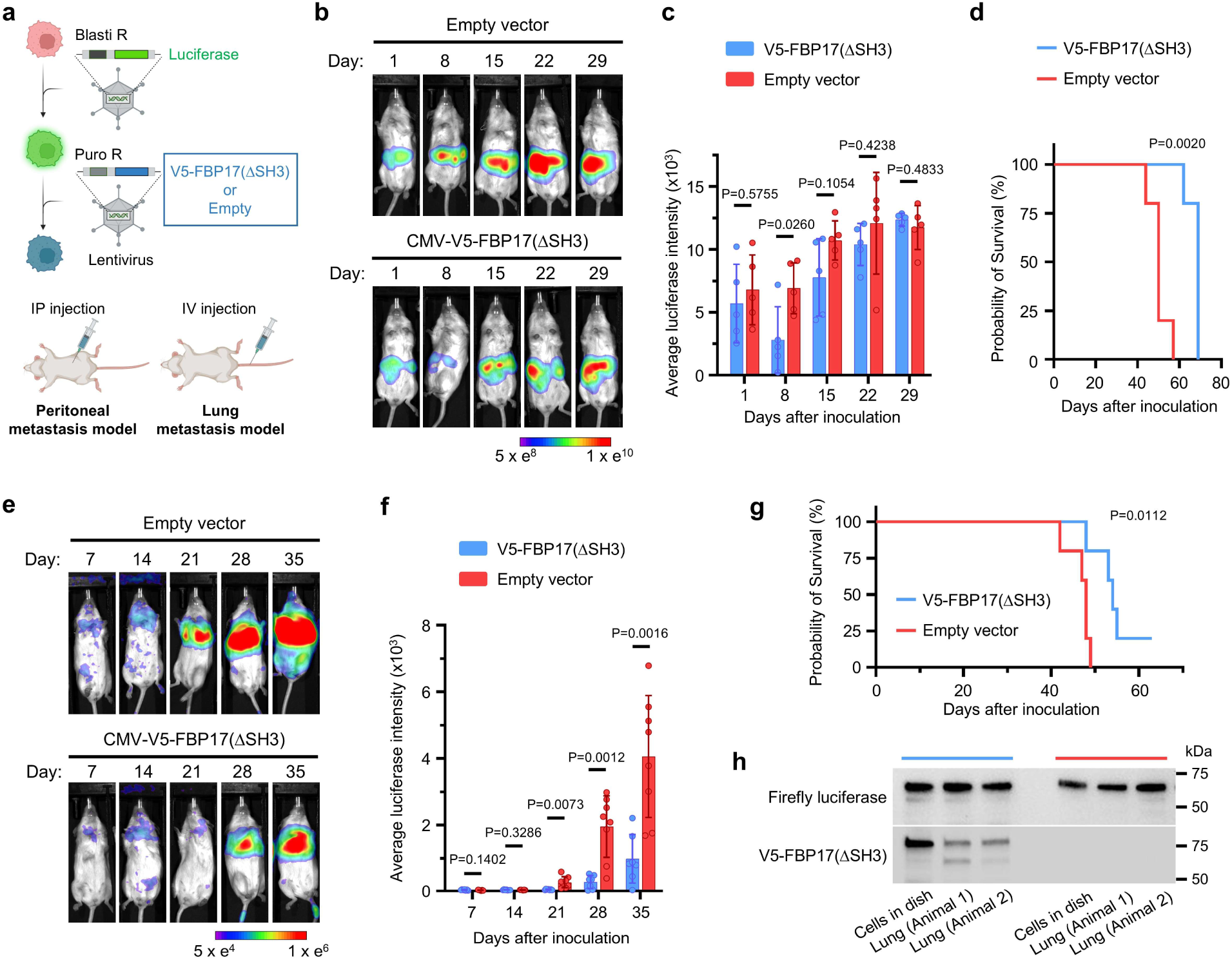
Blocking CIKA suppresses cancer metastasis in vivo. **a,** Schematic of xenograft models. Cancer cell lines were engineered by lentiviral transduction to stably express luciferase and either V5–FBP17(ΔSH3) or an empty-vector control. Left: BxPC3 cells were injected intraperitoneally to model peritoneal colonization. Right: MDA-MB-231 cells were injected via the tail vein to model hematogenous lung metastasis. **b**, Representative bioluminescent imaging of mice following intraperitoneal injection of BxPC3 cells. By day 8, V5–FBP17(ΔSH3) group showed markedly reduced signals compared to controls. See also Extended Data Fig. 6a for additional images. **c,** Quantification of bioluminescence signals after intraperitoneal injection. On day 8, V5–FBP17(ΔSH3) group exhibited ∼2.5-fold lower signals than controls and 2.0-fold lower relative to their own day-1 baseline, indicating impaired tumor colonization. *n* = 5 mice per group; mean ± SD; *p*-values from multiple unpaired t tests with Welch correction. **d,** Kaplan–Meier survival analysis showing significantly prolonged survival for V5–FBP17(ΔSH3) group (median = 69 days) compared with controls (median = 50 days). *p*-value from log-rank test. **e,** Representative bioluminescent images of mice after tail vein injection of MDA-MB-231 cells. By week 3, all control mice had developed detectable lung metastases, whereas the V5–FBP17(ΔSH3) group showed no detectable signal. Metastatic outgrowth in V5–FBP17(ΔSH3) group was delayed until weeks 4–5 and one V5–FBP17(ΔSH3) mouse remained tumor-free throughout the study. See also Extended Data Fig. 6b for additional images. **f,** Quantification of bioluminescence after tail vein injection confirming delayed metastatic progression in V5–FBP17(ΔSH3) group compared to controls. *n* = 7 [V5–FBP17(ΔSH3)] and 8 [control] mice; mean ± SD; *p*-values from multiple unpaired t tests with Welch correction. **g,** Kaplan–Meier curves showing prolonged survival in V5–FBP17(ΔSH3) group (median = 54 days, excluding the tumor-free mouse) compared with controls (median = 48 days). *p*-value from log-rank test. **h,** Western blot of lung tumor lysates compared with engineered cells. Tumor cells in mice show reduced V5–FBP17(ΔSH3) levels normalized to luciferase, suggesting selective outgrowth of cells with lower FBP17(ΔSH3) expression.

In the peritoneal colonization model, 3 × 10^5^ BxPC3 pancreatic cancer cells expressing Antares were injected intraperitoneally (IP) (**Fig. 6a**). Bioluminescent imaging one day post-injection showed comparable signals between FBP17(ΔSH3) and control groups, confirming that similar numbers of cells were initially delivered (**Fig. 6b** and **Extended Data Fig. 6a**). One week later, bioluminescence in the FBP17(ΔSH3) group was ∼2.5-fold lower than in controls and ∼2.0-fold reduced compared with its own day-1 baseline, indicating impaired tumor colonization (**Fig. 6b,c**). Although bioluminescence signals in both groups continued to rise and eventually plateaued, mice in the FBP17(ΔSH3) group survived significantly longer (median, 69 days) than controls (median, 50 days) (**Fig. 6d**). Therefore, blocking CIKA impairs early metastatic colonization and prolongs survival.

To evaluate the effect of CIKA inhibition on distant metastasis, we employed a lung metastasis model in which 1 × 10^5^ MDA-MB-231 cells expressing firefly luciferase were injected intravenously (IV) via the tail vein (**Fig. 6a**). Firefly luciferase was used because its longer imaging window after substrate administration facilitates detection of tumor growth in the lungs. Lung metastases became detectable by bioluminescence beginning at week 3. At this time, all eight control mice showed clear metastatic growth in the lungs, whereas none of the seven FBP17(ΔSH3) mice displayed detectable signals (**Fig. 6e**,**f** and **Extended Data Fig. 6b**). By week 4, lung metastases appeared in five of seven FBP17(ΔSH3) mice, increasing to six of seven by week 5. Notably, one FBP17(ΔSH3) mouse remained tumor-free and survived beyond the study endpoint. FBP17(ΔSH3) expression delayed metastatic onset and prolonged survival (median, 54 days excluding the tumor-free mouse) compared with controls (median, 48 days) (**Fig. 6g**).

To determine whether lung metastases in the FBP17(ΔSH3) group arose from cells with low FBP17(ΔSH3) expression, we performed western blot analysis (**Fig. 6h**). Both FBP17(ΔSH3) and firefly luciferase were robustly expressed in engineered MDA-MB-231 cells. However, in lung lysates containing metastatic lesions from two FBP17(ΔSH3) mice, FBP17(ΔSH3) levels were markedly lower than in the engineered cells when normalized to luciferase. These results suggest that the metastases originated from a subpopulation of cancer cells with initially low FBP17(ΔSH3) expression or from cells that downregulated FBP17(ΔSH3) during *in vivo* selection. Together, these findings demonstrate that blocking CIKA suppresses both local peritoneal colonization and distant lung metastasis.

## Discussion

In this study, we identified a mechanism termed curvature-induced kinase activation (CIKA), in which plasma-membrane curvature promotes the activation of oncogenic Src signaling by spatially organizing signaling molecules into dynamic, liquid-like condensates. Unlike receptor- or adhesion-mediated Src activation, CIKA arises from a biophysical cue, membrane geometry. Metastatic cells often exhibit reduced plasma-membrane tension accompanied by extensive membrane deformations, and recent studies have revealed strong associations between such membrane properties and the metastatic potential of solid tumors^19–23,26–30^. Our findings provide an additional mechanistic perspective on these associations, indicating that abnormal membrane mechanics in malignant cells can directly potentiate oncogenic signaling.

Curvature-sensing BAR-domain proteins act as adaptors that link membrane geometry to signaling. They oligomerize at curved membranes through their crescent-shaped BAR domains and recruit partners, often via their SH3 domains^31,51,52^. Our findings extend this mechanism by showing that curvature-induced oligomerization of TOCA-family proteins increases the local density of SH3 domains, driving biomolecular condensation through multivalent SH3-PRM interactions. These condensates function as signaling scaffolds that concentrate Src in its open conformation, enhance autophosphorylation at Y419, and exclude the inhibitory kinase Csk. Through this mechanism, geometrically defined membrane regions become spatially restricted kinase-activation hubs. Such curvature-induced condensation may represent a general strategy to enrich and compartmentalize biomolecules, ensuring the robustness and specificity of membrane-proximal signaling.

Beyond the Src/PI3K/Akt axis, curvature-induced condensates may scaffold additional signaling modules. Adaptor proteins involved in CIKA, such as cortactin and Tks5, participate in multiple signaling pathways, including Src/Abl, MAPK/ERK, and YAP/TAZ^94–97^. Recent studies have shown that membrane blebbing and curvature remodeling can activate PI3K/Akt and MAPK/ERK signaling, which are scaffolded by the septin cytoskeleton^28–30^. Although septins sense micrometer-scale curvature, distinct from the nanoscale curvature recognized by TOCA-family proteins in CIKA, additional work is needed to clarify how these systems intersect through shared downstream pathways or spatial coordination. Future studies should systematically map the signaling proteome of CIKA, delineate the hierarchy of associated pathways, and determine how CIKA hubs are regulated by mechanical cues or extracellular stimuli.

By using membrane curvature as an alternative organizing cue, metastatic cells bypass the requirement for focal adhesions and promote Src/PI3K/Akt signaling to resist anoikis. Because CIKA depends on nanoscale membrane architecture and multivalent scaffolding, rather than kinase overexpression or mutation, it represents a distinct form of oncogenic Src activation. Disrupting CIKA through a dominant-negative FBP17(ΔSH3) construct impairs anchorage-independent survival and metastatic colonization, highlighting the importance of this mechanism in metastasis. These findings suggest potential therapeutic strategies that target aberrant membrane deformation and curvature-induced condensates, rather than directly inhibiting kinase catalytic activity. Such approaches could target a unique biophysical and molecular vulnerability of metastatic cells to suppress tumor dissemination while sparing normal adherent cells. Developing drug-like molecules that can modulate CIKA will be crucial for turning this discovery into clinical treatments..

## Methods

### Nanofabrication and electron microscopy

Quartz wafers (Silicon Materials, 04Q 525-25-1F-SO) were spin-coated with CSAR 6200 and Electra 92 resists (AllResist), then patterned using JEOL JBX-6300FS e-beam lithography. A 120-nm-thick chromium mask was deposited via an AJA e-beam evaporator and the resists were lifted off in acetone/isopropanol. Patterns were transferred into vertical nanostructures by anisotropic reactive ion etching (C₄F₈/H₂/Ar, PT-Ox) for 3 min, and the chromium mask was removed with etchant 1020 (Transene). The resulting nanostructures were sputter-coated with a ∼3 nm platinum layer to improve conductivity and imaged by scanning electron microscopy (FEI Nova NanoSEM 450).

### Plasmid construction

DNA fragments encoding human SFKs (Src, Yes, Fyn, Fgr, Lck, Hck, Blk, Lyn and Frk), Csk, WIP, Pacsin2, FCHSD2, Tks4, Tks5, FAK, p130Cas and Sam68 were amplified from complementary DNA (cDNA) of U-2 OS or Jurkat cells. The C-terminal regulatory tyrosines of these SFKs [Src(Y530), Yes(Y537), Fyn(Y531), Fgr(Y523), Lck(Y505), Hck(Y522), Blk(Y501), Lyn(Y508) and Frk(Y497)] were mutated to phenylalanine by PCR using primers carrying the desired substitutions. Src truncation constructs—including Src(ΔSH4, Y530F, aa 21–536), Src(ΔSH3, Y530F, aa 1–83–GGSGG–146–536), Src(ΔSH2, Y530F, aa 1–150–GGSGG–249–536), Src(ΔSH1, aa 1–269), Src(SH3–SH2, aa 81–253), Src(SH3, aa 81–150) and Src(SH2, aa 142–253)—were generated by PCR or overlapping PCR. These DNA fragments were used to replace the EGFP in pEGFP-N1 for mammalian expression, or inserted into pEGFP-N1, pEGFP-C1, pmCherry-N1 or pmCherry-C1 vector (Clontech) for the expression of GFP- or RFP-tagged constructs.

N-WASP-GFP and GFP-TOCA-1 were gifts from David G. Drubin (UC Berkeley). The N-WASP coding sequence was amplified from N-WASP-GFP and used to replace the EGFP in pEGFP-N1 to express untagged N-WASP. GFP-Nck1 was a gift from Louise Larose (Addgene #45903). DNA fragments of p85ɑ and its tandem SH2 domain (p85α[nSH2–cSH2], aa 327–429) were amplified from pSV human p85α-HA (Addgene #11499, gift from Ronald Kahn), and inserted into pEGFP-C1 to express p85ɑ-GFP and p85ɑ(nSH2-cSH2)-GFP. GFP-cortactin was a gift from Anna Huttenlocher (Addgene #26722); the cortactin coding sequence was amplified and cloned into ptdTomato-N1 (Clontech) to create RFP-cortactin. PH-Akt-GFP was a gift from Tamas Balla (Addgene #51465).

GFP-FBP17 was a gift from Min Wu (Yale University). RFP-CIP4 was a gift from Christien Merrifield (Addgene #27685). The FCHo2 coding sequence was amplified from RFP-FCHo2 (Addgene #27686; gift from Christien Merrifield) and subcloned into pEGFP-N1 to produce GFP-FCHo2. Curvature-blocking mutants FBP17ΔSH3 (aa 1–549), CIP4ΔSH3 (aa 1–484) and TOCA1ΔSH3 (aa 1–479) were amplified and inserted into pEGFP-N1 and pmCherry-N1 to generate GFP- or RFP-tagged constructs. EGFP in pEGFP-C1 was replaced with EBFP2, and three SH3 domains of FBP17 (aa 548–617) or Nck1 (aa 1–61) were sequentially inserted to produce 3×SH3_FBP17_-BFP or 3×SH3_Nck1_-BFP (with 1×SH3_FBP17_-BFP as an intermediate). The pLenti pRRL-SV40(puro)_CMV vector was used as the empty control for lentiviral packaging. EBFP2, V5-FBP17ΔSH3 and BFP-FBP17ΔSH3 fragments were generated by PCR or overlap PCR and cloned into pLenti pRRL-SV40(puro)_CMV for lentivirus packaging. Lentiviral transfer vectors for Antares and Firefly luciferase were from the Michael Lin lab^98^.

### Cell culture and transient transfection

A549 (ATCC, CCL-185), U-2 OS (ATCC, HTB-96), HeLa (ATCC, CCL-2), MDA-MB-231 (ATCC, HTB-26), and BxPC3 (ATCC, CRL-1687) were cultured in cell culture medium, Dulbecco’s modified Eagle’s medium (DMEM) (Gibco) supplied with 10% (v/v) FBS (Sigma-Aldrich) and 1% (v/v) penicillin and streptomycin (Gibco). All cell lines were maintained at 37 °C in a 5% CO_2_ atmosphere.

A549 and U-2 OS cells were transiently transfected by electroporation. Briefly, cells cultured in 35-mm dishes were detached with TrypLE Express (Thermo Fisher Scientific), pelleted by centrifugation, and resuspended in a mixture of 100 µl electroporation buffer II (88 mM KH_2_PO_4_, 14 mM NaHCO_3_, pH 7.4), 2 µl electroporation buffer I (360 mM ATP, 600 mM MgCl_2_), and 0.1–2 µg total plasmid DNA. Electroporation was performed in 0.2-cm gap cuvettes using an Amaxa Nucleofector II (Lonza) according to the manufacturer’s protocol. Cells were immediately transferred into DMEM containing 10% (v/v) FBS and replated onto designated substrates. For transient expression of SFK constructs, 0.1 µg plasmid DNA was used and cells were cultured for 7 h before live-cell imaging or immunofluorescence to minimize the overexpression of active Src, which can cause cell detachment.

### Stable cell lines

Stable cell lines were generated by lentiviral transduction. For viral packaging, HEK293T cells (ATCC, CRL-3216) were co-transfected with lentiviral transfer plasmid together with packaging plasmid psPAX2 (Addgene #12260, gift from D. Trono) and envelope plasmid pMD2.G (Addgene #12259, gift from D. Trono) using Turbofect (Invitrogen) according to the manufacturer’s instructions. The transfection medium was replaced the next day with DMEM supplemented with 10% (v/v) FBS and 100 µM sodium pyruvate. Virus-containing supernatants were harvested 24 h post-transfection, clarified by centrifugation, and filtered through 0.45-µm PVDF syringe filters (Millipore). The viral supernatants were used to transduce cancer cell lines to stably express BFP, BFP–FBP17(ΔSH3), luciferases, or V5–FBP17(ΔSH3). Transduced cells were selected with 5 µg/ml blasticidin, 2.5 µg/ml puromycin, or both, and antibiotic selection was withdrawn 3 days prior to experiments.

### Xenograft models

All animal protocols were reviewed and approved by the Stanford Administrative Panel on Laboratory Animal Care (APLAC/IACUC). All procedures adhered to institutional and NIH guidelines, and all efforts were made to minimize animal use and suffering. NSG mice (Jackson Laboratory, RRID:IMSR_JAX:005557), 6–10 weeks old, were housed under pathogen-free conditions with a 12-h light/dark cycle and free access to food and water. Mice were randomly assigned to groups. For the peritoneal metastasis model, 3 × 10^5^ BxPC3 pancreatic cancer cells stably expressing NanoLuc and either V5–FBP17(ΔSH3) or empty control were injected intraperitoneally. For the lung colonization model, 1 × 10^5^ MDA-MB-231 breast cancer cells stably expressing Firefly luciferase and either V5–FBP17(ΔSH3) or empty control were injected intravenously via the tail vein. Tumor progression and metastatic burden were monitored at the indicated time points, by in vivo bioluminescence imaging using an IVIS Spectrum system (PerkinElmer). Mice were anesthetized with isoflurane and injected intraperitoneally with luciferase substrate D-luciferin (150 mg/kg) or fluorofurimazine (150 mg/kg) prior to imaging. Mice were anesthetized with ketamine and xylazine, and transcardially perfused with PBS followed to collect lungs for western blot analysis.

### Immunofluorescence labelling

For attached cells, cultures grown on nanostructured substrates were washed twice with PBS and fixed in 4% paraformaldehyde (Electron Microscopy Sciences) in PBS for 10 min at room temperature. Samples were rinsed three times with PBS, permeabilized with 0.1% Triton X-100 in PBS for 10 min, and blocked in 3% bovine serum albumin (BSA; Sigma) in PBS for 1 h at room temperature.

For suspended cells, cultures in tissue-culture-treated 6-well plates (∼50% confluency) were detached with TrypLE Express (Gibco) and gentle pipetting to obtain single cells. Cells were pelleted and resuspended in 500 µl DMEM containing 10% (v/v) FBS. Suspended cells were cultured at 37 °C, 5% CO_2_ for 15 min. Fixation was then performed by adding an equal volume of IC Fixation Buffer (Invitrogen) and incubating for 10 min at room temperature. Fixed cells were pelleted and washed three times with PBS by centrifugation, then resuspended in 30 µl collagen type I (2 mg ml⁻¹; Corning, 354236) and spread on glass-bottom dishes (Cellvis) for gelation. Collagen was neutralized with 10× PBS and kept at 4 °C until use. After 10 min gelation at 37 °C, collagen-embedded cells were permeabilized with 0.1% Triton X-100 in PBS for 20 min and washed three times with PBS (10 min each). Samples were then blocked in 3% BSA in PBS for 3 h at room temperature.

Following blocking, cells were incubated with primary antibodies diluted in 3% BSA/PBS overnight at 4 °C. After three PBS washes (10 min each), samples were incubated with dye-conjugated secondary antibodies or phalloidin diluted in 3% BSA/PBS for 3 h at room temperature, followed by three additional PBS washes (10 min each) before fluorescence imaging. Antibodies and dilutions are listed in **Supplementary Table 1**.

### Cell viability and colony formation assays

Single-cell suspensions were prepared using a method adapted from standard soft agar assays^88^. A549, MDA-MB-231, or BxPC3 cells stably expressing BFP–FBP17(ΔSH3) or BFP control were dissociated with TrypLE Express (Gibco) and gentle pipetting, pelleted, and resuspended in complete growth medium containing 0.35% low–melting-point agarose (Sigma) maintained at 37 °C. Cells (1 × 10⁴ per well in 200 µl) were plated into 24-well plates pre-coated with a 0.7% agarose base layer (200 µl) in complete medium. Each well was overlaid with 200 µl complete medium and cultured at 37 °C, 5% CO_2_.

For cell viability assays, cultures were stained using the LIVE/DEAD Viability/Cytotoxicity Kit (Invitrogen). Briefly, cells were incubated with 2 µM calcein-AM and 4 µM ethidium homodimer-1 in culture medium for 60 min at 37 °C. Viable cells (green) and dead cells (red) were visualized by epifluorescence microscopy using a ×10 objective, with z-stacks acquired through the agar top layer.

For soft agar colony formation assays, cultures were maintained for 14 days (A549, MDA-MB-231) or 21 days (BxPC3). The overlay medium was replaced with fresh DMEM containing 10% (v/v) FBS every 3 days. Colonies were imaged by bright-field microscopy using a ×2.5 objective with z-scanning through the agar top layer.

### Western blotting

Cells were lysed in ice-cold RIPA buffer (Thermo Fisher) supplemented with protease and phosphatase inhibitor cocktails (Thermo Fisher). Suspended cells were cultured at 37 °C, 5% CO_2_ for 90 min before lysis. For tumor lysates, snap-frozen lung tissues were minced on pre-chilled metal blocks and homogenized on ice with a micro-tip sonicator (1 s on/2 s off, 10–15 cycles) in an ice-cold RIPA buffer with protease and phosphatase inhibitors. Lysates were clarified by centrifugation (14,000 g, 20 min, 4 °C). Protein concentration was determined by BCA assay (Pierce). Samples were mixed with Laemmli buffer with 5% β-mercaptoethanol, heated at 95 °C for 5 min, and equal protein amounts were resolved on 4–15% TGX SDS–PAGE gels (Bio-Rad) and transferred to nitrocellulose membranes (Bio-Rad). Membranes were blocked in 5% non-fat milk in TBST (TBS + 0.1% Tween-20) for 1 h at room temperature.

Membranes were sectioned by molecular-weight range as needed and incubated with primary antibodies diluted in 5% BSA/TBST overnight at 4 °C. After three TBST washes (5–10 min each), membranes were incubated with HRP-conjugated secondary antibodies diluted in 5% BSA/TBST for 1 h at room temperature. Antibodies and dilutions used in Western blotting are listed in **Supplementary Table 1**. After three TBST washes, bands were developed using enhanced chemiluminescent substrate (SuperSignal West, Thermo Fisher) and imaged on a ChemiDoc MP (Bio-Rad).

### Epifluorescence, confocal imaging and FRAP

Epifluorescence images were acquired on a Leica DMi8 inverted microscope controlled by LAS X, equipped with a Leica LED8 excitation system and an sCMOS camera. Objectives used were 100× (1.40 NA oil), 63× (1.40 NA oil), 10× (0.30 NA air), and 2.5× (0.07 NA air). For live-cell imaging, cells were maintained in phenol-red–free DMEM (Gibco) with 10% (v/v) FBS at 37 °C and 5% CO_2_ in a stage-top incubator (Okolab). Z-stacks were acquired with step sizes of 10 µm for single-cell scans and 100 µm for colony scans through the agar top layer.

Confocal images were acquired on a Nikon A1R microscope controlled by NIS-Elements AR, using 405, 488, 561, and 633 nm laser lines. Confocal z-stacks were collected at 1.2 Airy Units using 20×/0.75 NA air or 60×/1.49 NA oil objectives. For FRAP, regions of interest (ROIs) were bleached for 1–2 bleach frames at 1 frame per second using the 488-nm laser line (∼10% power) alone for GFP or together with 561-nm laser line (∼10% power) for GFP and RFP, under the 60×/1.49 NA oil objective. Pre-bleach (5–10 frames) and post-bleach time series were acquired at 1 s or 5 s per frame. Live-cell conditions matched epifluorescence (phenol-red–free DMEM, 10% FBS, 37 °C, 5% CO₂, Okolab).

### Quantifications

All image processing used Fiji/ImageJ; custom MATLAB scripts with unified GUI were used to generate masks for each nanopillar and nanobar automatedly and place ROIs for intensity measurements; Cellpose v3^99^ were used and adapted to identify single cells and cell colonies.

### Nanobar end-to-side ratios

Masks for each nanobar were automatedly generated using MATLAB. Nanobar masks touching cell or image borders were excluded. Background was subtracted using an out-of-cell ROI. For each cell, masks were applied to compute an averaged nanobar image for each channel. End ROIs (circular, radius 11 pixel) and side ROIs (2-pixel-thick rectangular bands spanning the bar sides) were overlaid on the averaged image, and their mean intensities were measured. The end/side ratio from the averaged nanobar was reported as one data point per cell (*n* = cells). For gradient nanobars, nanobar images were averaged across the 40 bars of the same width within each array to yield one value per width per array (n = arrays). For thick nanobars (width 300-1000 nm), end ROIs are defined as half annuli (width 2 pixel).

### Nanopillar quantification

Analyses included nanopillar enrichment and two-channel correlation at nanopillars. Masks for each nanopillar were automatedly generated using MATLAB. Nanopillar masks touching cell or image borders were excluded. Background was subtracted using an out-of-cell ROI. For each cell, masks were applied to compute an averaged nanopillar image for each channel. To calculate nanopillar enrichment, an at-nanopillar circular ROI (radius 11 pixel) and a flat-surrounding concentric annulus ROI (width 12 pixel) were overlaid on the averaged image, and their mean intensities were measured; enrichment was calculated as the at-pillar/flat-surrounding ratio. For two-channel correlation, mean intensities from both channels within the at-pillar ROI were compared across nanopillars within a given cell to obtain the Spearman’s correlation coefficient. Each metric was reported as one data point per cell (*n* = cells).

### Ratiometric Src autophosphorylation analysis

Paired images of Src(pY419) and Src–GFP (or Src(Y530F)–GFP) were background-subtracted (rolling-ball radius 500 px), and pixel-wise ratiometric images were generated for analysis. For each cell, nanopillar masks as well as at-nanopillar and flat-surrounding ROIs were generated as described in “nanopillar quantification”. Fluorescence images of Src(Y530F)–GFP were auto-thresholded (Moments) in Fiji to generate binary masks; nanopillar regions were removed from these masks to yield focal-adhesion ROIs. Curvature-condensate ROIs were defined from the at-nanopillar ROIs with mean intensity values greater than 1.1, excluding nanopillars without condensates. The curvature-condensate, flat-surrounding, and focal-adhesion ROIs were overlaid on the ratiometric images and their mean values were measured, average per cell, and reported as one data point (*n* = cells). Pillar masks touching cell or image borders were excluded.

### FRAP analysis

Time series were background-subtracted (rolling-ball radius 100 px). Post-bleach time series were corrected for acquisition photobleaching using Fiji’s Bleach Correction (exponential fit). Intensities at the bleached nanopillars were measured from circular ROIs (radius 11 px) centered on the nanopillar, exported to GraphPad Prism 10, and normalized so that the pre-bleach mean = 1. With the first post-bleach point taken as I0, recovery curves were fit to a one-phase association model I(t) = I0 + (Plateau − I0) × (1 − e^−kt^); half-times were reported as t_1/2_ = ln(2)/k.

### Cell counting for viability assays

Live/dead fluorescence z-stacks through the agar top layer were maximum-intensity projected. The green (calcein-AM, live) and red (ethidium homodimer-1, dead) channels were processed independently in Fiji. Because cell density was low, segmentation was effectively performed using background subtraction (rolling-ball radius 50 px), auto-threshold (Triangle), and watershed separation. Objects were measured with Analyze Particles using a size filter (area > 25 µm²) and circularity 0.3–1.0 to exclude debris. Counts from both channels were exported and combined. Cell death rate was reported per well as % dead = #dead / (#live + #dead) (*n* = wells).

### Quantification of colony numbers

Bright-field z-stacks through the agar top layer were maximum-intensity projected. Colonies were segmented with Cellpose v3 using the pretrained the Segment Anything Model (SAM) as the starting point^99^; this model was fine-tuned on manually annotated images from this dataset and then applied to all images with fixed parameters. Resulting instance masks were imported into Fiji for measurement; to exclude single cells and debris, only objects with Feret diameter > 100 µm were counted as colonies. Counts were reported per well (*n* = wells).

### Statistics and reproducibility

All quantitative data are presented as mean ± SD. All in vitro experiments were independently repeated at least two times, unless noted otherwise in the legends, yielding consistent results. Statistical analyses were performed using GraphPad Prism (v10). Data distributions were assessed for normality using the Shapiro–Wilk test. For pairwise comparisons, two-tailed unpaired t-tests with Welch’s correction were used for parametric data, and the two-tailed Mann–Whitney U tests were used for non-parametric data. For datasets involving multiple pairwise comparisons, multiple t-tests with Welch’s correction were applied. Comparisons with a fixed value were analyzed using two-tailed one-sample t-tests. For comparisons involving more than two groups, one-way ANOVA followed by Dunnett’s T3 multiple comparisons was used for parametric data, and the Kruskal–Wallis test with Dunn’s multiple comparisons was used for non-parametric data. Kaplan–Meier survival curves were analyzed using the log-rank test. Exact *p*-values and sample sizes (*n*) are provided in the Figures and Figure legends. A *p*-value < 0.05 was considered statistically significant. No statistical method was used to predetermine sample sizes, which were chosen on the basis of previous studies in the field. The detailed test statistics, including degrees of freedom, are provided in **Supplementary Table 2**. Randomization and blinding were not applied. No data were excluded from the analyses, except for Fig. 6g, with the reason provided in the text.

## Reporting summary

Further information on research design is available in the Reporting Summary linked to this article.

## Data availability

Plasmids generated in this study, together with their sequence information, will be deposited in Addgene and distributed under materials transfer agreements (MTAs). All data are available in the main text and the supplementary information. Additional details and unique materials that support the findings of this study are available from the corresponding author upon reasonable request.

## Code availability

Custom MATLAB scripts used for data analysis will be made publicly available on GitHub. Additional details are available from the corresponding author upon reasonable request.

## Acknowledgments

We acknowledge Carolyn R. Bertozzi (Stanford University) for access to and support with the confocal microscopy. Nanofabrication was carried out with assistance from the Stanford Nanofabrication Facility and the Stanford Nano Shared Facilities. We thank David G. Drubin (UC Berkeley) for N-WASP-GFP and GFP-TOCA1; Louise Larose (Addgene #45903) for GFP-Nck1; Ronald Kahn (Addgene #11499) for p85α; Anna Huttenlocher (Addgene #26722) for GFP-cortactin; Tamas Balla (Addgene #51465) for PH-Akt-GFP; Min Wu (Yale University) for GFP-FBP17; and Christien Merrifield (Addgene #27685, #27686) for RFP-CIP4 and RFP-FCHo2.

This study was funded by the National Institutes of Health (R35GM141598, R01HL165491, R01NS121934, and R01CA311218), the Ono Pharma Breakthrough Initiative, and a Packard Fellowship for Science and Engineering awarded to B.C. Nanofabrication was carried out at the Stanford Nanofabrication Facility, a member site of the NSF National Nanotechnology Coordinated Infrastructure, and at the Stanford Nano Shared Facilities, supported by the National Science Foundation (award ECCS-2026822). C.-H.L. received additional support through a fellowship from the Stanford University Center for Molecular Analysis and Design.

## Author contributions

W.Z. and B.C. conceived the study. W.Z. designed the experiments, performed the majority of the research, and analyzed the data. X.Z., H.Y., and W.Z. conducted the animal experiments under the supervision of M.Z.L. and B.C. H.Y., C.-H.L., and A.A. contributed to plasmid construction. X.Y.Z. performed the SEM measurements. Z.J. contributed to nanostructure fabrication. W.Z. and B.C. wrote the manuscript with input from all authors. B.C. supervised the project.

## Ethics declarations

### Competing interests

The authors declare no competing interests.

### Inclusion & Ethics Statement

All authors meet the authorship criteria and contributed equitably to the study. This research did not involve global fieldwork or international human participant recruitment.

## Supplementary information

Supplemental Table 1: Antibodies used in this study. Supplementary Table 2: Test statistics

## Supplementary Videos

### Supplementary Video 1: FRAP of GFP–Nck1 in curvature-induced condensates at nanopillars

Live-cell FRAP showing rapid fluorescence recovery of GFP–Nck1, co-expressed with N-WASP and WIP, in curvature-induced condensates at nanopillars (t_1/2_ ∼5 s), demonstrating their liquid-like dynamics. Scale bar, 10 μm.

### Supplementary Video 2: FRAP of Src(Y530F)–GFP in curvature-induced condensates at nanopillars

Live-cell FRAP showing rapid fluorescence recovery of Src(Y530F)–GFP in curvature-induced condensates at nanopillars (t_1/2_ ∼32 s), confirming their dynamic molecular exchange and liquid-like behavior. Scale bar, 10 μm.

### Supplementary Video 3: FRAP, fusion, and fission of cytosolic condensates

Live-cell time-lapse imaging of cytosolic condensates formed by 3×SH3_FBP17_-BFP, GFP–Nck1, N-WASP, and WIP. GFP–Nck1 displayed rapid FRAP (two droplets in yellow box), and condensate droplets underwent fusion and fission events characteristic of liquid-liquid phase separation. Scale bar, 10 μm.

### Supplementary Video 4: FRAP of GFP–Nck1 and Src(ΔSH4, Y530F)–RFP in cytosolic condensates

Live-cell FRAP showing rapid fluorescence recovery of GFP–Nck1 and slightly slower recovery of Src(ΔSH4, Y530F)–RFP in cytosolic condensates formed together with cortactin, 3×SH3_FBP17_–BFP, N-WASP, and WIP, consistent with the FRAP dynamics of Nck1 (faster) and Src (slower) in curvature-induced condensates at nanopillars. Scale bar, 5 μm.

---

**Extended Data Fig. 1:**
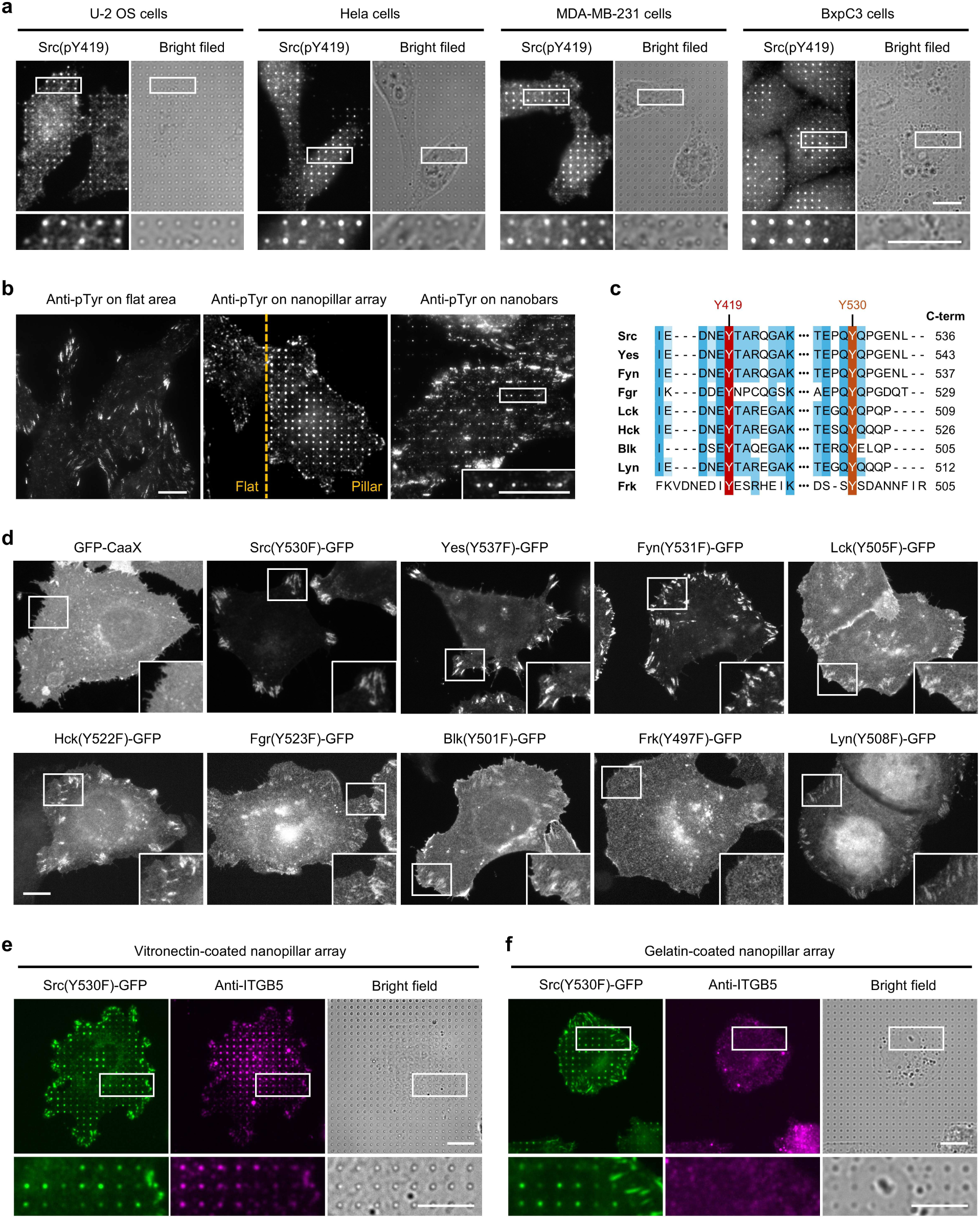
Additional characterization of active Src family kinase enrichment at membrane curvature. **a,** Src(pY419) enrichment at nanopillars is observed in multiple cell lines, including U-2 OS, HeLa, MDA-MB-231, and BxPC3, indicating that curvature-induced Src activation occurs broadly in different cell types. Scale bars, 10 µm. **b,** Pan-phosphotyrosine (pTyr) staining in U-2 OS cells shows that pTry primarily localizes to focal adhesions on flat regions. pTyr is also significantly enriched at curved membranes around nanopillars and nanobar ends. Scale bars, 10 µm. **c**, Sequence alignment of the activation loop surrounding Tyr419 across all nine human SFKs, showing high conservation of the phospho-epitope recognized by the Src(pY419) antibody. The conserved C-terminal autoinhibitory tyrosine residue (Y530 equivalent) is also shown for all nine SFKs. **d**, Subcellular localization of GFP-tagged open-conformation SFK mutants and GFP–CaaX in cells on flat regions. All SFK mutants except Frk localize to focal adhesions. Scale bar, 10 µm. **e,** Src(Y530F)–GFP localization relative to integrin β5 (ITGB5), a marker of curved adhesion, on vitronectin-coated nanopillars. Src(Y530F) does not colocalize with ITGB5, indicating that curvature enrichment of Src(Y530F) occurs independently of curved adhesions. Scale bars: 10 µm. **f**, Src(Y530F)–GFP is enriched at gelatin-coated nanopillars, which do not support curved adhesion formation (ITGB5 shows diffuse, uniform distribution). Results further confirm that curvature enrichment of active Src is independent of curved adhesions. Scale bars, 10 µm.

**Extended Data Fig. 2:**
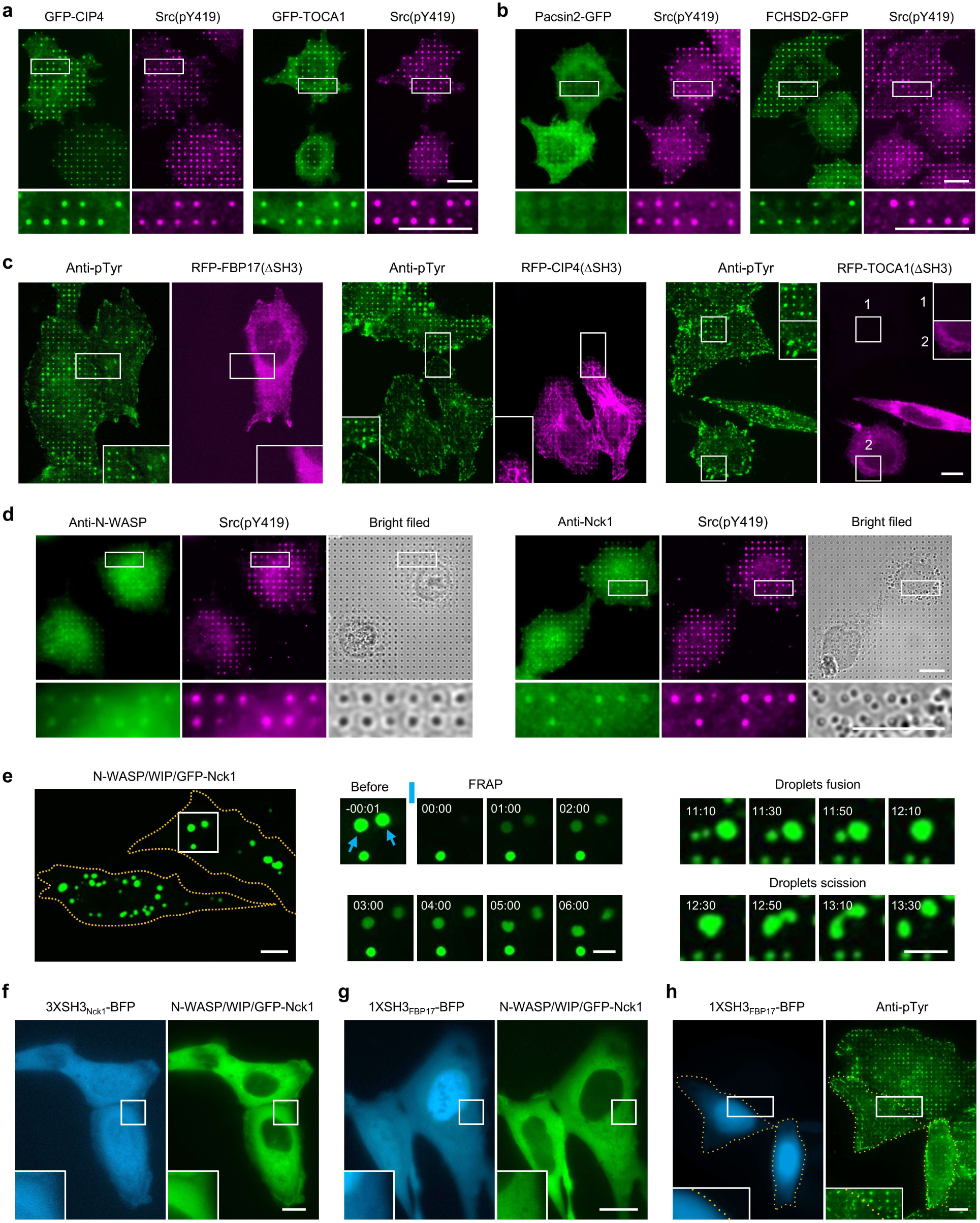
Additional evidence that membrane curvature–induced Src enrichment requires phase condensation. **a**, GFP–CIP4 and GFP–TOCA1 co-accumulate with Src(pY419) at nanopillars, similar to GFP–FBP17. Scale bars, 10 µm. **b,** Pacsin2–GFP does not accumulate at nanopillars. FCHSD2–GFP shows curvature enrichment but no correlation with Src(pY419). Scale bars, 10 µm. **c,** Expression of SH3-deletion mutants of FBP17, CIP4, or TOCA1 eliminates pTyr accumulation at nanopillars but preserves pTyr at focal adhesions. Scale bar, 10 µm. **d**, NCK and N-WASP, detected by immunofluorescence in untransfected cells, colocalize with Src(pY419) at nanopillars, confirming that curvature-induced condensation and Src enrichment correlate at the endogenous level. Scale bars, 10 µm. **e,** Cytosolic droplets formed by 3×SH3_FBP17_–BFP and N-WASP/WIP/GFP–Nck1 (left) showed rapid FRAP (middle) and dynamic fusion/fission (right), consistent with liquid-like dynamics. Scale bars, 10 µm; 5 µm (montages). See also Supplementary Video 3. **f**, A trivalent SH3 construct from Nck1 (3×SH3_Nck1_–BFP) fails to induce condensation of N-WASP/WIP/GFP–Nck1, indicating that SH3 multivalency alone is insufficient for condensate formation; TOCA-family SH3 domains are specifically required. Scale bar, 10 µm. **g,** A monovalent SH3 construct 1×SH3_FBP17_–BFP remains diffuse in the cytosol and does not induce condensation of N-WASP/WIP/GFP–Nck1. Scale bar, 10 µm. **h**, 1×SH3_FBP17_–BFP reduces pTyr accumulation at nanopillars but not at focal adhesions. Scale bar, 10 µm.

**Extended Data Fig. 3:**
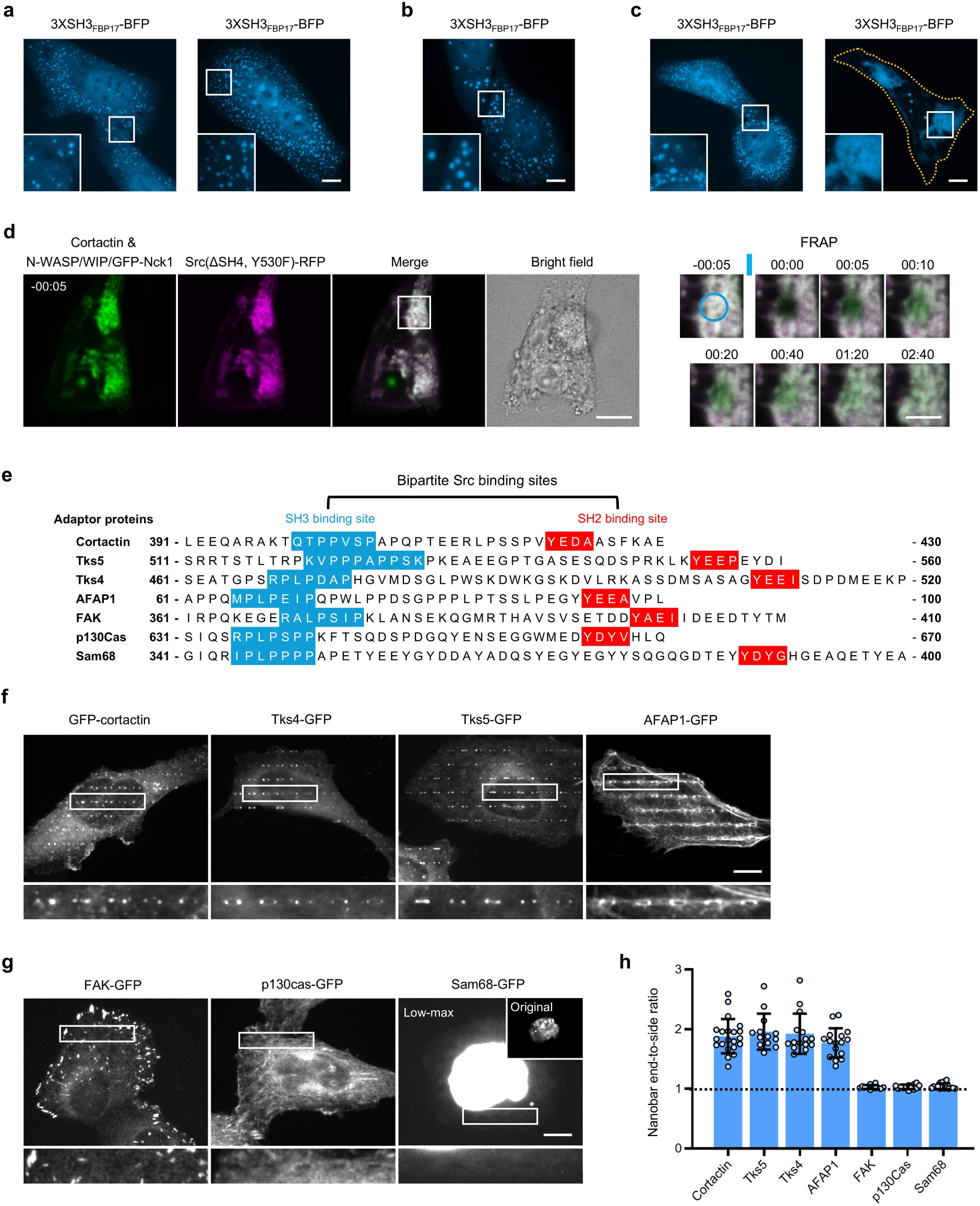
Additional evidence that curvature-induced condensates recruit Src via its SH3–SH2 domains and adaptor proteins. **a,** BFP channel of cytosolic condensates formed by 3×SH3_FBP17_–BFP, GFP–Nck1, N-WASP, and WIP, which fail to recruit Src(SH3–SH2) or Src(ΔSH4, Y530F). Scale bar, 10 µm. Related to Fig. 3d. **b,** BFP channel of cytosolic condensates formed by 3×SH3_FBP17_–BFP, GFP–Nck1, N-WASP, and WIP, which recruit cortactin. Scale bar, 10 µm. Related to Fig. 3f. **c,** BFP channel of cytosolic condensates formed by cortactin together with 3×SH3_FBP17_–BFP, GFP–Nck1, N-WASP, and WIP, which recruit Src(SH3–SH2) or Src(ΔSH4, Y530F). Scale bar, 10 µm. Related to Fig. 3g. **d,** FRAP analysis of GFP–Nck1 and Src(ΔSH4, Y530F)–RFP in cytosolic condensates reveals rapid fluorescence recovery, indicating liquid-like dynamics. Nck1 recovers slightly faster than Src(ΔSH4, Y530F). Scale bars, 10 µm; 5 µm (montages). See also Supplementary Video 4. **e,** Sequences of the SH2- and SH3-binding motifs in adaptor proteins known to engage Src in a bipartite manner. Adaptor proteins include cortactin, Tks5, Tks4, AFAP1, FAK, Sam68, and p130Cas. **f** and **g,** Representative images of GFP-tagged adaptor proteins on nanobar substrates. Cortactin, Tks4, Tks5, and AFAP1 accumulate at nanobar ends, whereas FAK, Sam68, and p130Cas do not. Scale bars, 10 µm. **h**, Quantification of curvature enrichment (nanobar end-to-side ratios) for adaptor proteins shown in (e-g). Cortactin, Tks4, Tks5, and AFAP1 show significant enrichment, while FAK, Sam68, and p130Cas do not. *n* = 12 cells per condition, from two independent experiments; data are mean ± SD.

**Extended Data Fig. 4:**
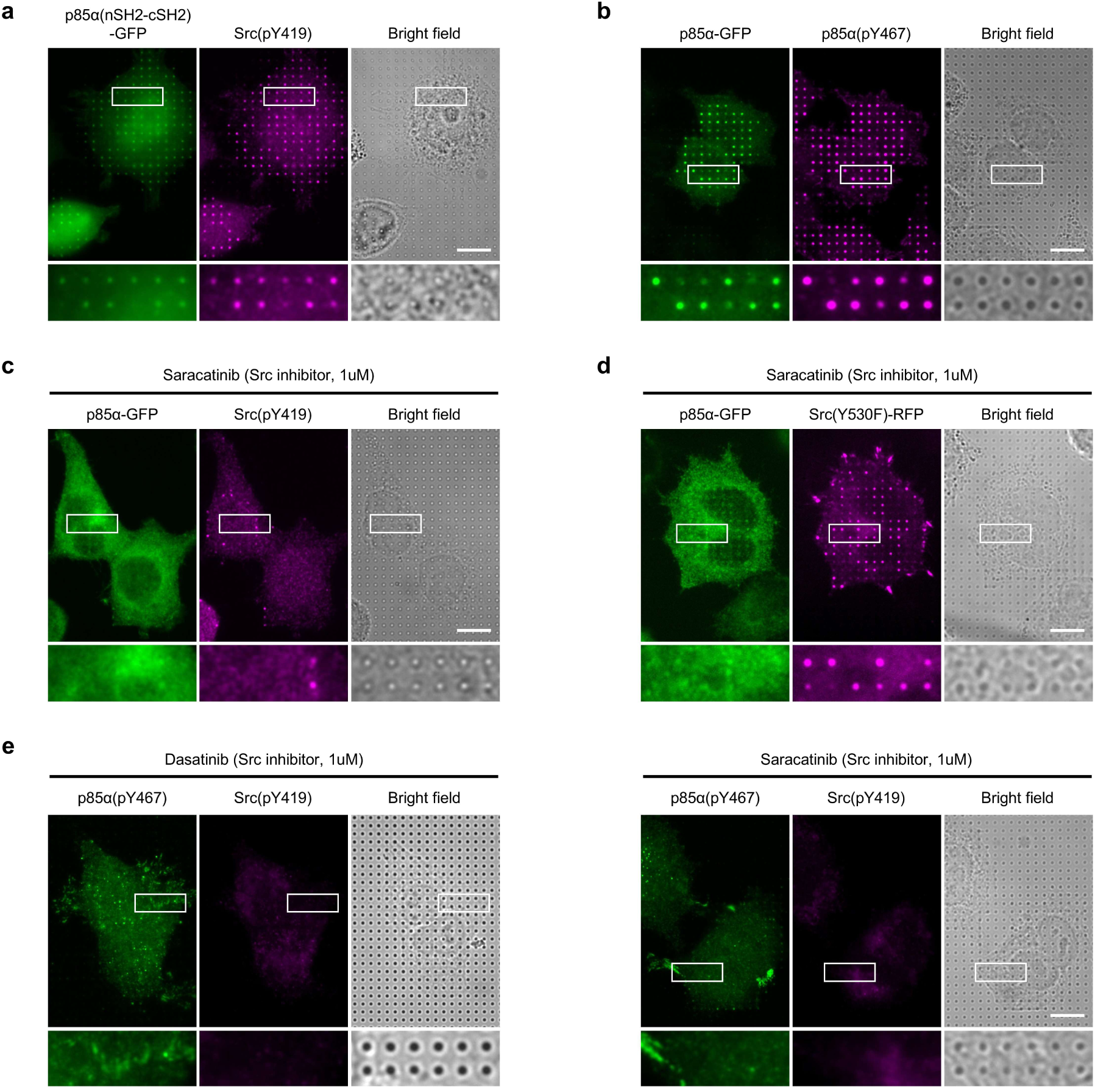
Additional evidence that curvature-induced Src activation promotes PI3K/Akt signaling. **a**, GFP-tagged nSH2–cSH2 fragment of p85α accumulates at nanopillars and colocalizes with Src(pY419). Scale bar, 10 µm. **b,** Immunostaining of p85α(pY467) showing its strong curvature-dependent enrichment at nanopillars, colocalizing with Src(pY419). Scale bar, 10 µm. **c**, Treatment with Src inhibitor saracatinib abolishes the enrichment of p85α–GFP and Src(pY419) at nanopillars, similar to the effect of dasatinib. Scale bar, 10 µm. **d,** Despite saracatinib treatment, Src(Y530F)–RFP remains enriched at nanopillars, while p85α–GFP enrichment is abolished, indicating that Src kinase activity is necessary for PI3K but not Src recruitment to curvature. Scale bar, 10 µm. **e,** Immunostaining confirms that both dasatinib and saracatinib eliminate nanopillar enrichment of Src(pY419) and p85α(pY467). Scale bar, 10 µm.

**Extended Data Fig. 5:**
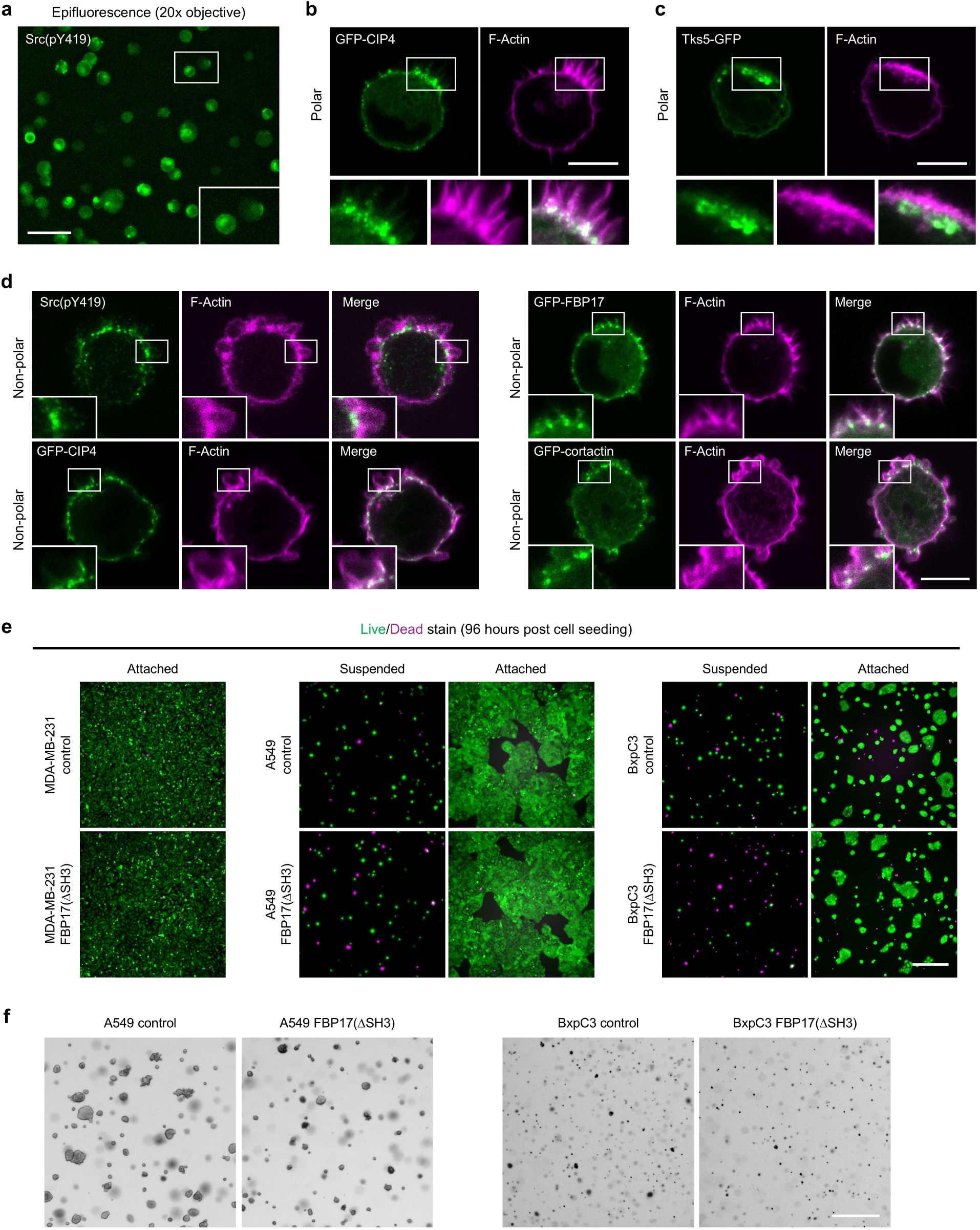
Additional evidence for curvature-induced Src signaling in suspended cancer cells. **a,** Large-field epifluorescence image showing that Src (pY419) is not uniformly distributed on the cell surface. Scale bar, 50 µm. **b** and **c,** 2D confocal images showing that GFP–CIP4 (B) and Tks5–GFP (C) preferentially localize to crown-base invaginations rather than crown tips in suspended A549 cells. Scale bar, 10 µm. **d,** In non-polarized suspended cells lacking a dominant crown, Src(pY419), GFP–FBP17, GFP–CIP4, and GFP–cortactin are enriched at membrane invaginations distributed along the cortex. Scale bar, 10 µm. **e,** Live/dead staining showing that FBP17(ΔSH3) expression reduces survival of A549 and BxPC3 cells suspended in soft agar, consistent with results from MDA-MB-231 cells in Fig. 5i and quantification in Fig. 5j. Scale bar, 500 µm. **f,** Representative images of soft-agar colony formation assays showing that FBP17(ΔSH3)-expressing A549 and BxPC3 cells formed markedly fewer large colonies compared to controls, consistent with MDA-MB-231 cells in Fig. 5l and quantification in Fig. 5m. Scale bar, 500 µm.

**Extended Data Fig. 6:**
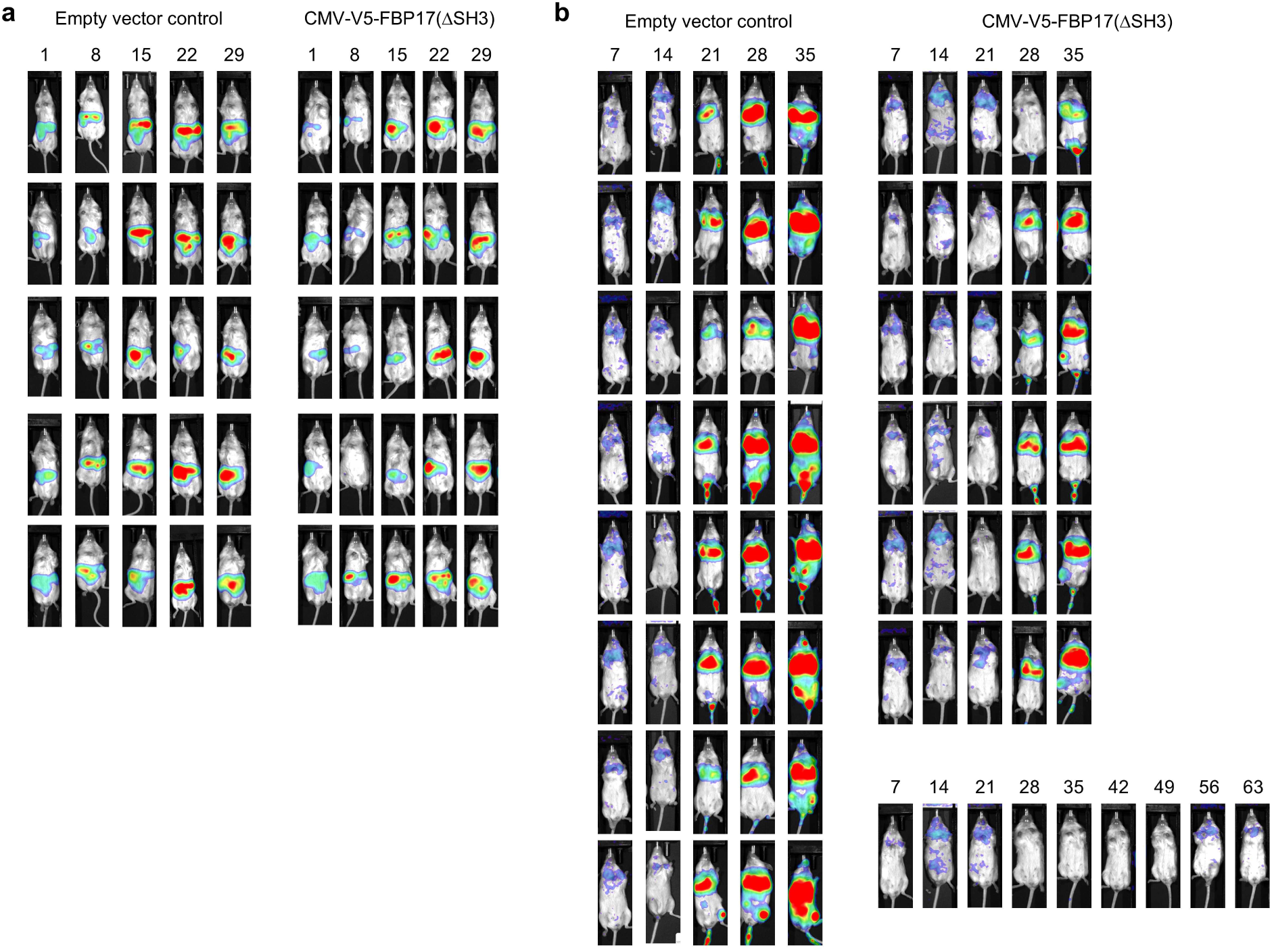
Full bioluminescence imaging series for xenograft models. **a**, Complete bioluminescence imaging series for all mice (*n* = 5 per group) injected intraperitoneally with BxPC3 cells and imaged from day 1 to day 29. Images correspond to the representatives shown in Fig. 6b. **b**, Complete bioluminescence imaging series for all mice injected via tail vein with MDA-MB-231 cells. n = 7 [V5–FBP17(ΔSH3)] and 8 [control] mice, imaged weekly from day 7 to day 35. Images correspond to the representatives shown in Fig. 6e.

## Notes

### Competing Interest Statement

The authors have declared no competing interest.

